# Direct linkage detection with multimodal IVA fusion reveals markers of age, sex, cognition, and schizophrenia in large neuroimaging studies

**DOI:** 10.1101/2021.12.13.472507

**Authors:** Rogers F. Silva, Eswar Damaraju, Xinhui Li, Peter Kochunov, Aysenil Belger, Judith M. Ford, Daniel H. Mathalon, Bryon A. Mueller, Steven G. Potkin, Adrian Preda, Jessica A. Turner, Theo G.M. van Erp, Tulay Adali, Vince D. Calhoun

## Abstract

With the increasing availability of large-scale multimodal neuroimaging datasets, it is necessary to develop data fusion methods which can extract cross-modal features. A general framework, multidataset independent subspace analysis (MISA), has been developed to encompass multiple blind source separation approaches and identify linked cross-modal sources in multiple datasets. In this work we utilized the multimodal independent vector analysis model in MISA to directly identify meaningful linked features across three neuroimaging modalities — structural magnetic resonance imaging (MRI), resting state functional MRI and diffusion MRI — in two large independent datasets, one comprising of control subjects and the other including patients with schizophrenia. Results show several linked subject profiles (the sources/components) that capture age-associated decline, schizophrenia-related biomarkers, sex effects, and cognitive performance. For sources associated with age, both shared and modality-specific brain-age deltas were evaluated for association with non-imaging variables. In addition, each set of linked sources reveals a corresponding set of multi-tissue spatial patterns that can be studied jointly.

## 1. Introduction

Multimodal neuroimaging data can provide rich information to better understand brain structures and functions, and boost biomarker detection (Uludag and Roebroeck, 2014). Although analysis of each data modality separately can yield important insights into the structural or functional integrity of the brain, the relationship between different views from multimodal neuroimaging data is often complex and unknown. Data-driven approaches are ideal for such cases, leveraging naturally occurring associations across modalities to discover structure-function relationships, which may not occur in the same regions and may covary among subjects in complex ways. As more research institutions participate in open science data sharing practices, large-scale neuroimaging datasets (>1000 subjects) with multimodal data are becoming widely available, giving researchers opportunities to develop novel approaches to multimodal fusion analysis that can offer insights into cross-modal (joint) associations and identify important missing links in brain development and mental disorders (Calhoun and Sui, 2016).

Blind source separation (BSS) techniques, in particular independent component analysis (ICA) (Comon, 1994; Bell and Sejnowski, 1995), have gained popularity in neuroimaging analysis because they make minimal assumptions about the latent sources, are readily available, and yield interpretable results. However, ICA is only suitable for single modality data. Several other methods have been developed to extract multimodal features, including canonical correlation analysis (Hotelling, 1936), extensions of ICA such as joint ICA (Calhoun et al., 2006) and parallel ICA (Liu et al., 2008), as well as independent vector analysis (IVA) (Adali et al., 2014; Kim et al., 2006), which generalizes ICA to multiple datasets. Recently, we proposed a data-driven blind source separation model called multidataset independent subspace analysis (MISA) (Silva et al., 2021) that generalizes many basic BSS techniques, such as ICA and IVA, to recover *subspaces* (i.e., a collection of linked latent sources) within and across multiple datasets simultaneously. MISA utilizes the Kotz distribution (Kotz, 1975) to model source distributions, thus leveraging all-order statistics (second- and higher-order) to model the underlying latent subspaces. Another advantage of MISA is that it allows comparison across many different types of BSS because they are special cases of the general model it implements. In this work, we use MISA to implement the multimodal IVA (MMIVA) model described in Section 2.4 and demonstrate its utility on two large neuroimaging datasets: one including typical older adults from the UK Biobank enhanced imaging study, and the other including pooled data from multiple studies/sites that investigate psychosis and age-matched controls.

We evaluated MMIVA on standard derivative data from three neuroimaging modalities: 1) gray matter tissue probability segmentation (GM) maps from structural MRI (sMRI) data, which markedly convey regional variations in gray matter concentration with age, sex and other neurobiological factors, 2) amplitude of low frequency fluctuations (ALFF) maps computed from resting state functional MRI (rfMRI) scans, which inform about the strength of local connectivity as well as potential for long-range associations, and 3) fractional anisotropy (FA) maps obtained from diffusion MRI (dMRI) scans, which characterize the degree of directional water diffusion in white matter bundles. In the two independent datasets we considered, the results show several cross-modal associations are present. Moreover, we observed covariation of these linked sources with factors such as age, sex, and cognition, as well as group label (patients with psychosis vs control subjects).

In the following, Section II describes the data, preprocessing, and methodology utilized in this work. Section III presents our results, which are further discussed in Section IV before presenting our final conclusions.

## 2. Methods

### 2.1. Imaging Data

We used two large independent multi-site datasets to evaluate the MMIVA model. For the first dataset, we utilized imaging data from a subset of 3497 subjects participating in the UK Biobank study (Miller et al., 2016), a prospective epidemiological study with a large imaging database. Specifically, we utilized multivariate features (Calhoun and Adali, 2008) extracted from each subject and each data modality. All data included in our analysis were collected in two of the three participating locations in the United Kingdom. All participants provided informed consent from their respective institutional review boards.

T1-weighted structural MRI images were acquired using a 3D MPRAGE sequence at 1mm^3^ isotropic sagittal slices with acquisition parameters: 208 × 256 × 256 matrix, R=2, TI/TR=880/2000 ms. Diffusion MRI images were acquired using a standard Stejskal-Tanner spin-echo sequence at 2mm^3^ isotropic resolution at three different b values (b = 0, 1,000 and 2,000 s/mm^2^) and 50 distinct diffusion-encoding directions, each using a multi-band (MB) factor of 3. Resting functional MRI were acquired axially at 2.4mm^3^ resolution while the subjects fixated at a cross with the following acquisition parameters: 88 × 88 × 64 matrix, TE/TR=39/735ms, MB=8, R=1, flip angle 52°. A pair of spin echo scans of opposite phase encoding direction in the same imaging resolution as rfMRI scans were acquired to estimate and correct distortions in rfMRI echo-planar images, and a single-band high resolution reference image was acquired at the start of the rfMRI scan to ensure good realignment and normalization.

The second dataset includes pooled data from 4 studies that collected imaging data from patients with schizophrenia, schizo-affective disorder, bipolar disorder and age-matched healthy controls. These studies are Center of Biomedical Research Excellence in brain function and mental illness (COBRE) (Aine et al., 2017), function biomedical informatics network (fBIRN) (Keator et al., 2016), Maryland Psychiatric Research Center (MPRC), and Bipolar and Schizophrenia Network for Intermediate Phenotypes (BSNIP) (Tamminga et al., 2014). The dataset information is summarized in Table 1.

**Table 1:** Dataset demographics information. Demographics in the UK Biobank dataset and the patient datasets are shown in the table below. (*N*_*M*_ : number of male subjects; *N*_*F*_ : number of female subjects; *N*_*C*_: number of controls; *N*_*P*_ : number of patients.)

| Dataset | Site | $N$ | $N_M$ | $N_F$ | $N_C$ | $N_P$ | Mean Age (std) (y) | Age Range (y) |
| --- | --- | --- | --- | --- | --- | --- | --- | --- |
| UK Biobank Dataset |  |  |  |  |  |  |  |  |
| UKB | 1 | 2379 | 1201 | 1178 | N/A | N/A | 63.1 (7.2) | 48-79 |
| UKB | 2 | 528 | 251 | 277 | N/A | N/A | 61.8 (7.3) | 46-79 |
| Patient Datasets |  |  |  |  |  |  |  |  |
| COBRE | 1 | 160 | 123 | 37 | 85 | 75 | 37.79 (12.8) | 18-65 |
| MPRC | 1 | 60 | 44 | 16 | 26 | 34 | 38.67 (13.2) | 16-62 |
| MPRC | 2 | 106 | 43 | 63 | 62 | 44 | 40.28 (13.1) | 16-64 |
| MPRC | 3 | 1 | 1 | 0 | 0 | 1 | 55.00 (0.0) | 55-55 |
| BSNIP | 1 | 237 | 133 | 104 | 137 | 100 | 37.62 (14.7) | 16-65 |
| BSNIP | 2 | 192 | 103 | 89 | 99 | 93 | 40.13 (13.2) | 15-65 |
| FBIRN | 1 | 24 | 19 | 5 | 16 | 8 | 36.00 (9.9) | 22-51 |
| FBIRN | 2 | 17 | 14 | 3 | 8 | 9 | 40.88 (9.9) | 23-57 |
| FBIRN | 3 | 53 | 41 | 12 | 27 | 26 | 42.81 (12.5) | 19-60 |
| FBIRN | 4 | 54 | 44 | 10 | 27 | 27 | 36.04 (11.4) | 20-58 |
| FBIRN | 5 | 16 | 9 | 7 | 7 | 9 | 38.62 (9.4) | 22-53 |
| FBIRN | 6 | 44 | 21 | 23 | 24 | 20 | 34.93 (10.6) | 19-58 |
| FBIRN | 7 | 35 | 30 | 5 | 20 | 15 | 37.91 (10.2) | 18-60 |

### 2.2. Subject Measures Data

UK Biobank provides extensive phenotype information for each subject, including age, sex, lifestyle measures, cognitive scores, etc. We used a subset of the subject measures (SM) reported in (Miller et al., 2016) to identify associations between the subject demographics and the multivariate source component vectors (SCVs) obtained from MMIVA. Following the approach in Smith et al. (2015), we dropped subjects with more than 4% missing data. This resulted in *N* = 2907 (out of 3497) subject scores for MANCOVAN analysis (see Section 2.5). Of 64 SMs, we dropped 10 columns which had extreme values. Extreme values are identified in 2 steps. First, the sum of squared absolute median deviations (ssqamdn) for each SM is computed. If there is any SM with max(ssqamdn) > 100 × mean(ssqamdn), then that SM has subjects with extreme outliers which can influence statistical analysis and, thus, that SM is dropped. This resulted in 54 phenotypes including age, sex, fluid intelligence, a set of measures covering amount and duration of physical activity, frequency of alcohol intake, cognitive test scores, time spent watching TV, and sleep duration (see Miller et al. (2016) for details). For the measures that were retained, any missing values were imputed utilizing the K-Nearest Neighborhood method implemented in MATLAB’s knnimpute() (Cunningham and Delany, 2021).

The patient data SMs include only age, sex, and diagnosis information.

### 2.3. Preprocessing

We processed each of the three imaging data modalities to obtain GM, ALFF, and FA feature maps, which were then used for multimodal fusion analysis.

Specifically, the sMRI images underwent segmentation and normalization to MNI space using the SPM12 toolbox, yielding GM, white matter (WM), and cerebro-spinal fluid (CSF) tissue probability maps. The normalized GM segmentations were spatially smoothed using a 10mm FWHM Gaussian filter. The smoothed images were resampled to 3mm^3^. We defined a group mask to restrict the analysis to GM voxels as follows. First, an average GM segmentation map from all subjects was obtained from the normalized GM segmentation maps at 1mm^3^ resolution. This map was binarized at a 0.2 threshold (classify voxels with a group average probability > 0.2 as gray matter) and resampled to 3 mm^3^ resolution, which resulted in *V*_*m*_ = 44,318 in-brain voxels.

We used distortion corrected, FIX-denoised (Griffanti et al., 2014), normalized rfMRI data provided by the UK Biobank data resource to compute subject-specific ALFF maps, defined as the area under the low frequency band [0.01-0.08 Hz] power spectrum of each voxel time course in a given scan. We then obtained mean-scaled ALFF maps (mALFF), which are subject-specific ALFF maps divided by their global mean ALFF value, since this scaling has been shown to result in greater test-retest reliability of ALFF maps (Zhao et al., 2018). The mALFF maps were smoothed using a 6mm FWHM Gaussian filter and resampled to 3mm isotropic voxels. We used the same group mask learned from GM features for mALFF maps in the subsequent fusion analysis.

For dMRI data, we used the FA maps provided by the UK Biobank consortium. The preprocessing steps that raw dMRI images underwent are thoroughly described in (Alfaro-Almagro et al., 2018). The FA maps were then spatially smoothed using a 6mm FWHM Gaussian filter and resampled to 3mm^3^ voxels. For dMRI data, we computed a group mask similar to the approach described above for sMRI data. However, the group average WM segmentation was binarized at a threshold of 0.4, resulting in *V*_*m*_ = 18,684 in-brain voxels in the group mask.

Prior to running our fusion model, we verified that none of the included voxels contained invalid values, such as NaN or +*/*−Inf. For each modality, we also performed variance normalization of the subject-specific spatial maps (mean removed, then divided by standard deviation), followed by mean removal per voxel (across subjects). Finally, site effects were regressed out voxelwise from the datasets to account for mean differences in feature maps due to scanner effects as follows:

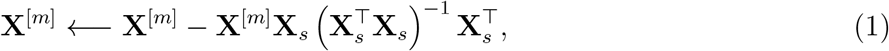

where **X**^[*m*]^ is the *V*_*m*_ × *N* data matrix from each modality, **X**_*s*_ = [**1 𝓁**_*s*_], with **1** being a *N* × 1 column vector of ones and **𝓁**_*s*_ being a column vector containing the site labels.

### 2.4. Multimodal IVA (MMIVA) fusion model

Here we present a general IVA approach for direct analysis of heterogeneous multimodal data. As mentioned earlier, independent vector analysis is a natural extension of independent component analysis. While ICA operates on a single dataset to obtain statistically independent source signals via estimation of *one* linear unmixing matrix, IVA performs joint estimation of *many* unmixing matrices simultaneously across multiple datasets (Kim et al., 2006).

Briefly, ICA is a blind source separation model that assumes linear mixing of *C* statistically independent sources **s**, yielding the observed data **x**:

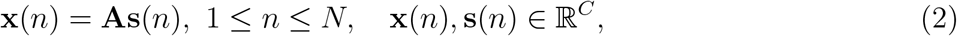

where **A** is the invertible mixing matrix, and *N* is the number of observations (here, the number of subjects). The ICA algorithm seeks to identify the sources **ŝ**(*n*) = **Wx**(*n*) by estimating an unmixing matrix **W**, according to certain properties of the sources such as higher-order statistics and non-Gaussianity. Another typical strategy for solving the ICA problem involves minimizing the mutual information among the sources:

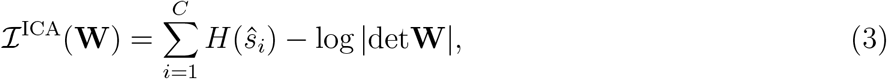

where *H*(*ŝ*_*i*_) is the differential entropy of the *i*-th source, given by 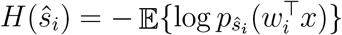.

IVA extends the ICA model to multiple (*M*) datasets, assuming a linear mixture of *C* independent sources for each dataset:

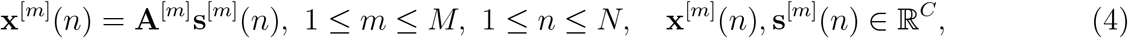

additionally assuming statistical *dependence* (i.e., linkage) among corresponding sources, i.e., those with same index *i*. Each *i*-th collection of linked sources is called a source component vector (SCV) and defined as 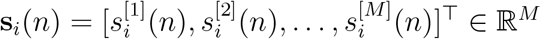. Here, *M* = 3, such that each SCV spans across the three modalities.

Solving the IVA problem involves minimizing the following mutual information:

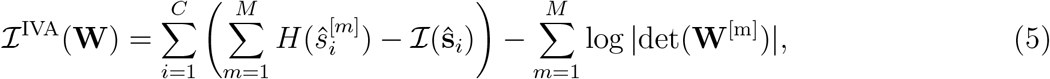

where the terms in the big parentheses correspond to the joint entropy of the *i*-th SCV, *H*(**ŝ**_*i*_), showing that IVA will seek independence *among* SCVs while capturing multimodal dependence *within* SCVs; the mutual information ℐ(**ŝ**_*i*_) indicates dependence among sources in the *i*-th SCV (Adali et al., 2014).

Here, we utilize the flexible MISA implementation (Silva et al., 2021) to estimate the multimodal IVA fusion model described above, leveraging both second- *and* higher-order statistics. This implementation enables direct data fusion even when **A**^[*m*]^ is a tall matrix and has a *different* number of rows (the modality’s intrinsic dimensionality) in each modality (Silva et al., 2014). The utility of this “transposed IVA” approach was also demonstrated in the sample-poor (low *N*) regime using only second-order statistics Adali et al. (2015b,a). A broader discussion with comparisons to other BSS approaches in data fusion is also available (Silva and Plis, 2019).

As depicted in Figure 1, we performed MMIVA fusion of the GM, mALFF, and FA features by treating each modality as one of the *M* datasets in the IVA model described above. In order to initialize **W**^[*m*]^, the multimodal data matrices **X**^[*m*]^ were reduced to 30 principal directions using multimodal group principal component analysis (MGPCA). Unlike standard PCA that finds orthogonal directions of maximal variation for each modality separately, MGPCA finds directions of maximal *common* variation, i.e., eigenvectors are computed based on the average of the *scaled* covariance matrices **Σ**^[*m*]^:

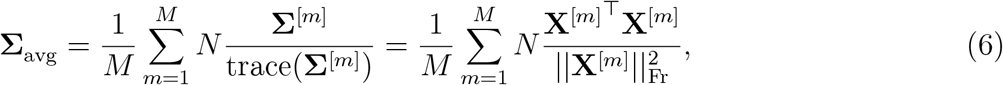

where|| · ||_Fr_ indicates the Frobenius norm. The scaling factor we used is 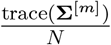, which is the ratio of the variance in the modality to the number of observations (subjects). Letting **Λ** and **H** be the top 30 eigenvalues (with largest absolute value) and eigenvectors of **Σ**_avg_, respectively, we define the reduced joint dataset **X**_*r*_ and corresponding whitening matrices **whtM**^[*m*]^ as follows:

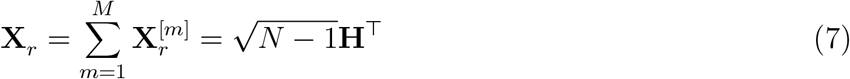

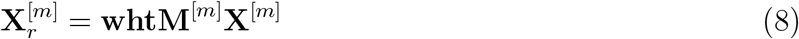

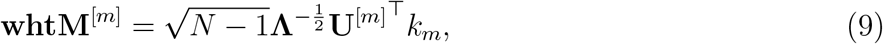

where 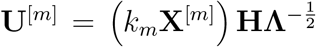, and 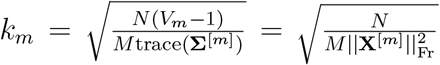. Following, the MGPCA-reduced data **X**_*r*_ underwent an ICA estimation using the Infomax objective (Bell and Sejnowski, 1995) to obtain 30 *common* independent sources **ŝ**_*I*_(*n*) = **W**_*I*_**x**_*r*_(*n*).

**Figure 1:**
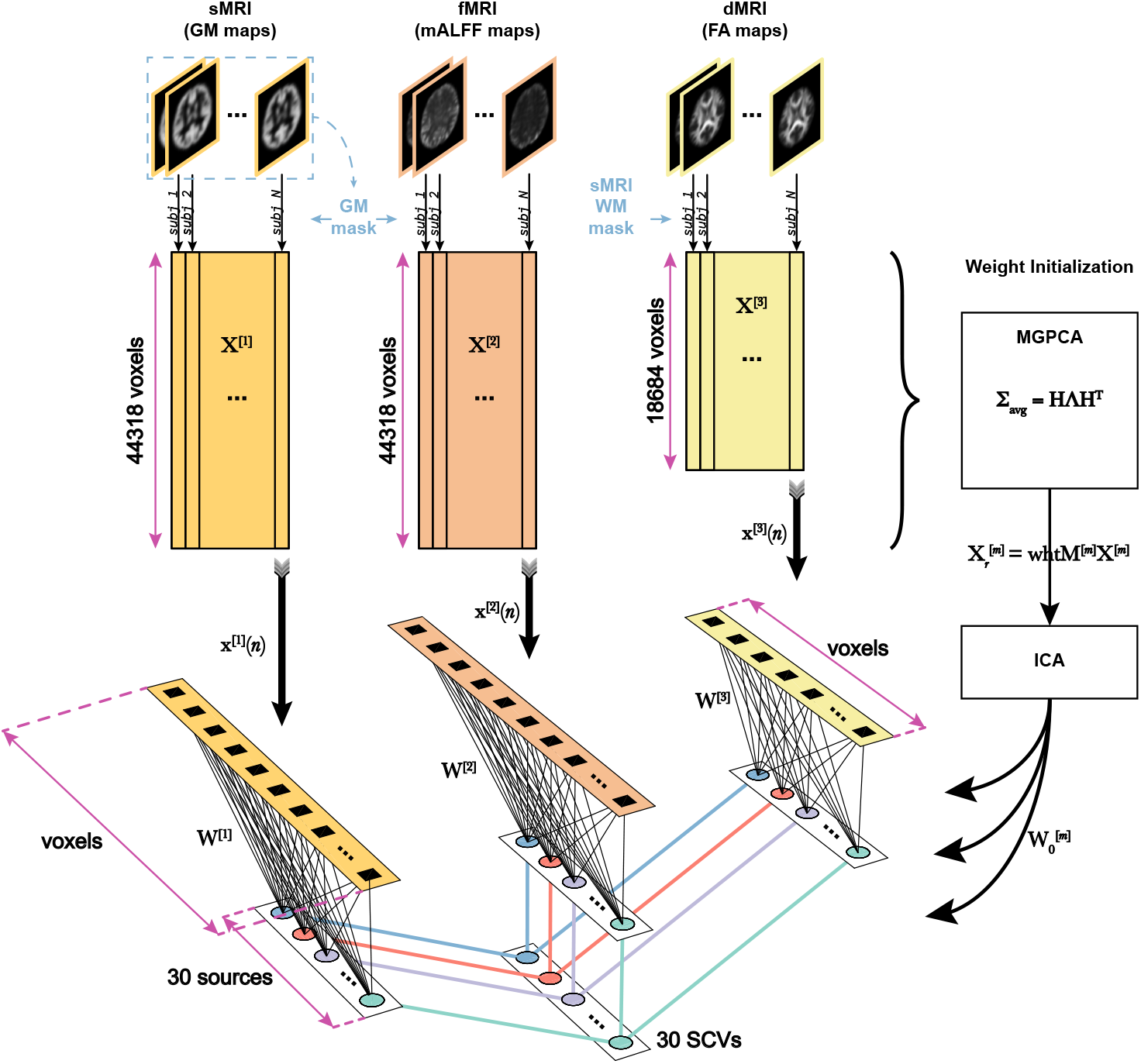
Multimodal IVA (MMIVA) fusion.

We improved upon the Infomax estimate by configuring and running MISA as an ICA model initialized with **W**_*I*_, but now assuming source distributions to follow a *univariate* Kotz distribution with parameters *λ* = 0.8966, *β* = 0.5462, *η* = 1, thus yielding **ŝ**_*K*_(*n*) = **W**_*K*_**x**_*r*_(*n*). The final combined MGPCA+ICA estimate 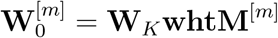 was utilized to initialize the MMIVA model.

MMIVA was then estimated directly from the full data **X**^[*m*]^ by simply reconfiguring and running MISA as an IVA model, yielding our final joint decompositions **ŝ**^[*m*]^(*n*) = **W**^[*m*]^**x**^[*m*]^(*n*). Spatial maps were then estimated using least squares as **Â** ^[*m*]^ = **X**^[*m*]^**Ŝ**^[*m*]T^(**Ŝ**^[*m*]^**Ŝ**^[*m*]T^)^−1^, where **Ŝ**^[*m*]^ is the *C* × *N* source matrix from each modality.

As discussed earlier, MMIVA accounts for dependence among corresponding sources across modalities. For both MISA and MMIVA models, the SCV distributions are assumed to take a multivariate Kotz distribution (Nadarajah, 2003; Kotz, 1974):

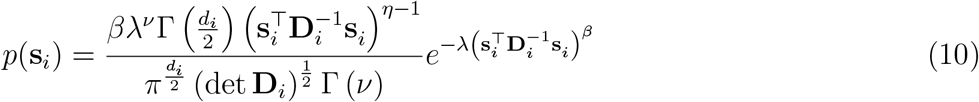

where *d*_*i*_ = *M* is the SCV dimensionality, *β >* 0 controls the shape of the distribution, *λ >* 0 controls the kurtosis, and 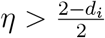 controls the hole size, while 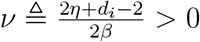 and 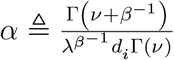 are defined for brevity. Γ (·) denotes the gamma function. The positive definite dispersion matrix **D**_*i*_ is related to the SCV covariance matrix 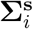 by 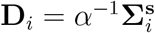. The multivariate Kotz distribution has been shown to generalize well across multivariate Gaussian, multivariate Laplace, and multivariate power exponential distributions (Anderson et al., 2013). For MMIVA, the Kotz parameters were set to *λ* = 0.8966, *β* = 0.5462, *η* = 1. The MISA methods were implemented using the MISA toolbox (Silva et al., 2021).

### 2.5. Statistics

We used the MANCOVAN toolbox to identify associations between the subject measures (SM) and the multivariate source component vectors (SCVs) obtained from MMIVA. The MANCOVAN Toolbox evaluates multivariate MANCOVA models and implements multivariate stepwise regression (backward selection mode) to identify associations between SM (predictors) and MMIVA sources (multivariate response) for each modality separately (one MANCOVA model per modality, each with *C* = 30 sources/responses). The stepwise regression approach eliminated insignificant SM terms at the *α >* 0.01 level at each step using the multivariate Lawley-Hotelling trace test. Subsequently, univariate tests with the surviving SM terms are reported for each of the 30 sources within each modality. Initial model selection based on multivariate tests significantly reduces the total number of tests performed. The univariate tests were performed and corrected for multiple comparisons at Bonferroni threshold (0.05/30, for 30 sources).

In addition to the SMs, the following nuisance covariates were included prior to stepwise regression:

- sMRI: correlation of warped subject GM segmentation map to mean GM segmentation map,
- dMRI: correlation of warped subject FA map to mean FA map,
- rs-fMRI: correlation of warped subject mALFF map to mean mALFF map, and mean framewise displacement (mFD) computed from rigid body movement estimates from resting fMRI scan realignment step.

Any variables with fewer than 8 levels were modeled as categorical variables and the rest were modeled as continuous variables. Only age by sex interaction was considered.

Eta-squared (*η*^2^) was used to measure the strength of the relationship between the predictors and the SCVs. *η*^2^ is defined as the ratio of variance explained (the sums-of-squares, or *SS*) in the dependent variable by a predictor while controlling for other predictors:

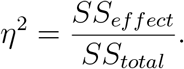

In the implementation, we calculated the unbiased estimator of the population’s *η*^2^, epsilon-squared (*ϵ*^2^). The reason we chose epsilon-squared (*ϵ*^2^) is that it has been found to be less biased compared to omega-squared (*ω*^2^) (Carroll and Nordholm, 1975). Detailed explanation of effect size indices can be found in Albers and Lakens (2018). Both Type II and Type III sum-of-squares are used to evaluate effect sizes. In general, we prioritize Type II statistics because Type II is more statistically powerful than Type III when no interactions are present (Langsrud, 2003). For those SCVs where an interaction term has significant *p*-value (*p <* 0.05*/*number of surviving SM), we replaced Type II by Type III statistics. We then identified SCVs with effect sizes *ϵ*^2^ > 0.02 (Cohen (1992)) for each predictor in each modality. SCVs are selected if the same SM meets this effect size criterion on at least two of the three modalities. The univariate tests are reported alongside effect sizes for completeness, but do not play a role in SCV selection.

### 2.6. Brain-age delta modeling on UK Biobank data

To further evaluate the significant age-related UK Biobank SCVs identified with MANCOVA, we evaluated the difference between the predicted brain age and the chronological age (the brain-age delta) in two steps, as described in Smith et al. (2020). In a brain age model, there are two main components: chronological age, which carries a normative essence to it, and deviations^1^. Then, building a model that approximates brain age will partially (since it’s an approximation) capture variability from each of the two components. Our interest is then on the physiological effects that explain the deviations from the expected (chronological) age. In that sense, when predicting brain age from imaging features, the interesting portion of the captured variability is that which is associated with the deviations because we want to understand what (non-imaging) factors associate with these deviations.

First, we conducted a partialling step in order to determine each source’s unique contribution. To that end, we regressed out all other sources from each source^2,3^. Inspection of these partialled sources revealed that partialled sMRI and dMRI features were highly anti-correlated. Thus, we chose to include only the partialled sMRI feature in this brain-age analysis (an analysis including partialled dMRI instead of sMRI is included in the supplementary section). We also noticed that, given the high correlation within SCVs, partialling by-and-large removed the shared (and the largest) portion of the variability within SCVs. Thus, we evaluated the top singular vector of each SCV to capture the shared information, based on the singular value decomposition (SVD) of the 3-column matrix containing all three sources from the same SCV, each normalized to have unit standard deviation. These shared-SVD features are naturally independent of each other because the SCVs from which they were derived are independent from each other to begin with. Since partialled terms have the shared information removed, the shared-SVD features are also at most only weakly correlated with partiallized terms. We verified that the correlation among the SVD-shared features and partialled sMRI, fMRI, and dMRI sources is low. Therefore, we selected the SVD-shared features, partialled sMRI, and partialled fMRI expression levels for the following brain-age analysis. We included an analysis with SVD-shared features, partialled dMRI, and partialled fMRI expression levels in the supplementary section 6.3.

In the first step of brain-age delta modeling, the subject expression levels (the top SVD-shared feature plus the partialled sMRI and fMRI features) from five SCVs with significant age effect size (based on the multivariate MANCOVA assessment in 2.5) were used to predict age. Thus, the initial estimate of brain-age delta was evaluated as:

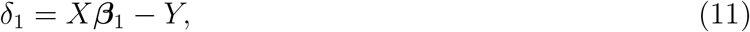

where *Y* is the actual age after removing the mean age across subjects, *X* is the matrix of significant age-related SCVs with size 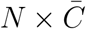 (*N* = number of subjects, 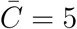 SVD-shared features + 5 sMRI sources + 5 dMRI sources = 15), and ***β***_1_ is the 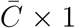 vector of regression parameters. As such, *δ*_1_ shows the aging effects that cannot be captured by the imaging modalities, with positive values suggesting accelerated aging and negative values suggesting slowed down aging.

In the second step of brain-age delta modeling, the model is refined to identify aspects of *δ*_1_ that cannot be captured by age terms (notice that *δ*_1_ is by definition dependent on age) nor confounds, such as sex. Thus, *δ*_2_ is evaluated as:

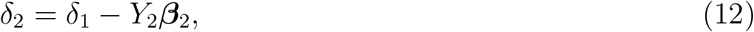

where the regression matrix *Y*_2_ includes the demeaned linear age term, the demeaned quadratic age term after regressing out the linear age effects and normalizing to have the same standard deviation as the linear age term, the demeaned cubic age term after regressing out the linear and quadratic age effects and normalizing to have the same standard deviation as the linear age term, sex, the interaction between sex and each of the three age terms, the framewise displacement variable, and the spatial normalization variables from three modalities. Note that after each effect removal step, the mean of the adjusted variable was removed again. ***β*** regression coefficients are estimated as usual by ***β***_1_ = (*X*^T^*X*)^−1^*X*^T^*Y* and ***β***_2_ = (*Y*_2_^T^*Y*_2_)^−1^*Y*_2_^T^*δ*_1_.

Following that, source-specific contributions to the overall brain age delta can be determined, yielding individual delta vectors, *δ*_1*i*_ and *δ*_2*i*_, as follows. First, note that the source-specific contributions can be combined to recover the original Y (here, 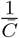 is arbitrary and irrelevant since age effects are removed in the following step, thus we set 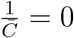:

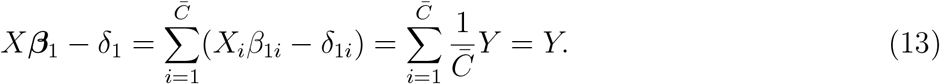

The source-specific delta vector at the first stage, *δ*_1*i*_, can be calculated by:

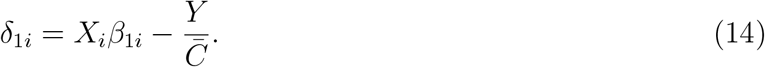

In the next step, we compute the regression coefficients ***β***_2*i*_ = (*Y*_2_^T^*Y*_2_)^−1^*Y*_2_^T^*δ*_1*i*_ and the individual delta vector at the second stage is obtained:

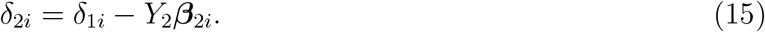

We removed the mean of the *δ*_2*i*_ at the end. We repeated the above individual brain-age delta estimation process (14) and (15) for each source in each age-related SCV.

We then computed partialled *δ*_2*i*_ by regressing out other *δ*_2*i*_ vectors from each particular *δ*_2*i*_ vector. Then we evaluated the Pearson correlations between subject measures and *δ*_2*i*_, as well as subject measures and partialled *δ*_2*i*_.

## 3. Results

Following SCV estimation by the proposed multimodal IVA approach, the MANCOVA procedures revealed significant predictors for each modality, including age, psychosis, sex, and prospective memory to name a few. Then, for each modality, we fit a univariate multiple regression model between each individual source in that modality and all significant predictors in order to measure effect sizes.

For the UK Biobank dataset, after MANCOVA stepwise regression, we fit linear models using the surviving variables as follows:

1. 9 variables (sex, time spent watching tv, age first had sexual intercourse, time to answer, fluid intelligence, age, physical exercise principal component 1, spatial normalization, interaction between sex and age) to predict each sMRI source;
2. 11 variables (alcohol, sex, sleep duration, age first had sexual intercourse, time to answer, fluid intelligence, age, physical exercise principal component 3, spatial normalization, mean framewise displacement, interaction between sex and age) to predict each fMRI source;
3. 8 variables (sex, time spent watching TV, time to answer, fluid intelligence, mean time to correctly identify matches, age, spatial normalization, interaction between sex and age) to predict each dMRI source.

The Type II effect size for the *age* * *sex* interaction term was lower than 0.02 for all UKB univariate linear models. Therefore, we only report Type II *ϵ*^2^ for the UKB dataset.

For the patient dataset, we fit linear models using the same 7 predictors (age, sex, diagnosis, the interaction between age and sex, the interaction between age and diagnosis, the interaction between sex and diagnosis, the interaction among age, sex and diagnosis) to predict unimodal sources for each of the three modalities. The Type II effect sizes of interaction terms are all small (*ϵ*^2^ *<* 0.02) so results from Type II model are reported.

The effect size measure indicated five age-related SCVs (3, 5, 8, 16, 17), five sex-related SCVs (9, 10, 16, 17, 22) and one SCV (22) associated with time to answer in the UKB dataset. In the patient dataset, two schizophrenia-related SCVs (14, 19) and six age-related SCVs (2, 4, 6, 18, 19, 20) were identified.

Next, we report the cross-modal Pearson correlations captured within the aforementioned SCVs. These reflect the degree of similarity between patterns of expression level across subjects. Table 2 below summarizes the strength of multimodal pairwise links recovered by MMIVA, indicating high similarity in expression levels between modalities, particularly between fMRI and dMRI. This is indicative of strong multimodal linkages (i.e., for the reported SCVs, the corresponding spatial patterns of each modality are expressed at largely similar levels in each subject).

**Table 2:** Cross-modal Pearson correlations within SCVs.

| SCV | sMRI-fMRI | fMRI-dMRI | sMRI-dMRI |
| --- | --- | --- | --- |
| SCVs in UK Biobank dataset |  |  |  |
| 3 | 0.720 | 0.840 | 0.667 |
| 5 | 0.639 | 0.764 | 0.613 |
| 8 | 0.643 | 0.920 | 0.634 |
| 9 | 0.692 | 0.873 | 0.663 |
| 10 | 0.740 | 0.700 | 0.703 |
| 16 | 0.773 | 0.908 | 0.746 |
| 17 | 0.728 | 0.785 | 0.693 |
| 22 | 0.642 | 0.539 | 0.692 |
| SCVs in patient datasets |  |  |  |
| 2 | 0.705 | 0.634 | 0.603 |
| 4 | 0.649 | -0.666 | -0.637 |
| 6 | 0.758 | 0.807 | 0.709 |
| 14 | 0.623 | 0.699 | 0.549 |
| 18 | 0.7 | 0.722 | 0.595 |
| 19 | 0.742 | 0.884 | 0.768 |
| 20 | 0.67 | 0.788 | 0.675 |

Following, we show Pearson spatial map correlations between the two datasets for each modality. In Figure 2, the SCVs from the patient dataset were sorted from highest to lowest correlation in the sMRI modality since that modality showed the largest evidence of cross-dataset similarities. Notice the age-related SCV 8 in the UK Biobank dataset is highly correlated with age- and schizophrenia-related SCV 19 in the patient dataset, in both sMRI and dMRI modalities, indicating replication of the multimodal link identified by MMIVA. Detailed descriptions of UK Biobank SCV 8 and patient SCV 19 are presented in Sections 3.1 and 3.2, respectively.

**Figure 2:**
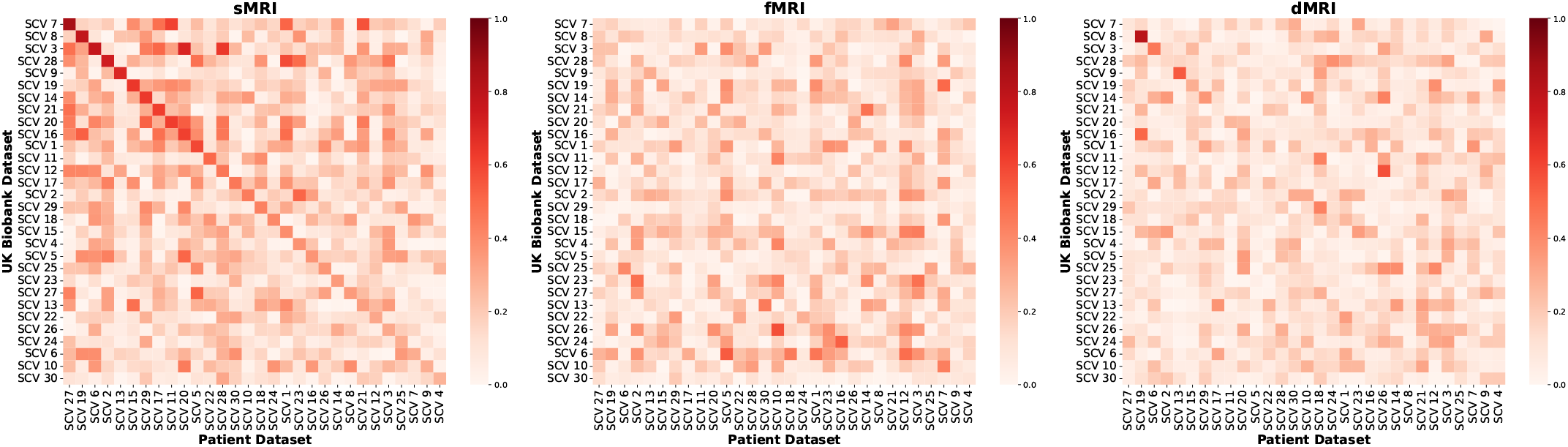
Pearson spatial map correlations between the two datasets for each modality. Absolute values of Pearson correlations between spatial maps from the two datasets were evaluated for each modality. Both datasets were processed separately, with no cross-contamination. The sMRI correlations showed the largest evidence of cross-dataset similarities. Thus, SCVs were matched across datasets based on sMRI correlations using the Jonker-Volgenant algorithm for the linear sum assignment problem. The best-matched sMRI correlations were then sorted in descending order. The resulting optimal permutation of patient SCVs was then applied to the other modalities.

### 3.1. Associations with aging in UK Biobank

For the UK Biobank data, five SCVs (3, 5, 8, 16 and 17) showed age effects consistently for all three modalities. The sources from all SCVs with age effects had their sign oriented in the direction of decline with age. The 3D scatter-plot of linked subject expression profiles (sources) in SCV 8, which is the SCV carrying the most significant age association, is presented in Figure 3. Consistent with Table 2, a strong linear association can be observed between modalities. The corresponding mixing weights (spatial maps) are also presented and depict in hot colors the regions expressing larger values at younger age (likewise, expressing lower values at older age) for each of the three modalities. In sMRI (gray matter), age-associated *reductions* were observed in the dorsal and ventral caudate, nucleus accumbens, rostral and caudal temporal thalamus, caudal hippocampus, ventro- and dorso-medial parieto-occipital sulci, several areas of the anterior and posterior cingulate gyrus, and several areas by the sylvian fissure (mainly in the superior temporal and dorsal insular gyri), with age-related *increases* in sensorimotor areas, visuomotor area 7, lateral occipital gyrus, as well as the inferior frontal junction and parts of the nucleus accumbens and putamen. Similar levels of subject-specific age-associated *reductions* were observed in fMRI (ALFF), in areas such as the dorsal prefrontal cortex, Broca’s area, the caudal angular and supramarginal gyri, the extreme part of inferior temporal gyrus adjacent to the fusiform gyrus, and portions of the visual cortex including the cuneus, the ventro-medial part of the parieto-occipital sulcus, and the superior part of the occipital polar cortex, with age-related *increases* in sensorimotor, thalamus, and parietal/occipital areas including the angular gyrus, visuomotor area 7, the lateral superior occipital gyrus, the middle occipital gyrus, and the inferior part of the occipital polar cortex. In dMRI (FA), airalogous subject-specific age-related *reductions* occurred in the anterior portion of the corpus callosum, forceps minor, superior longitudinal fasciculus II, the anterior thalamic radiation, and the optic radiation, with age-related *increases* in the corticospinal tract, superior thalamic radiation, caudal parts of the arcuate fasciculus, and the middle longitudinal fasciculus. The remaining age-associated sources are shown in the supplementary section Figure 9.

**Figure 3:**
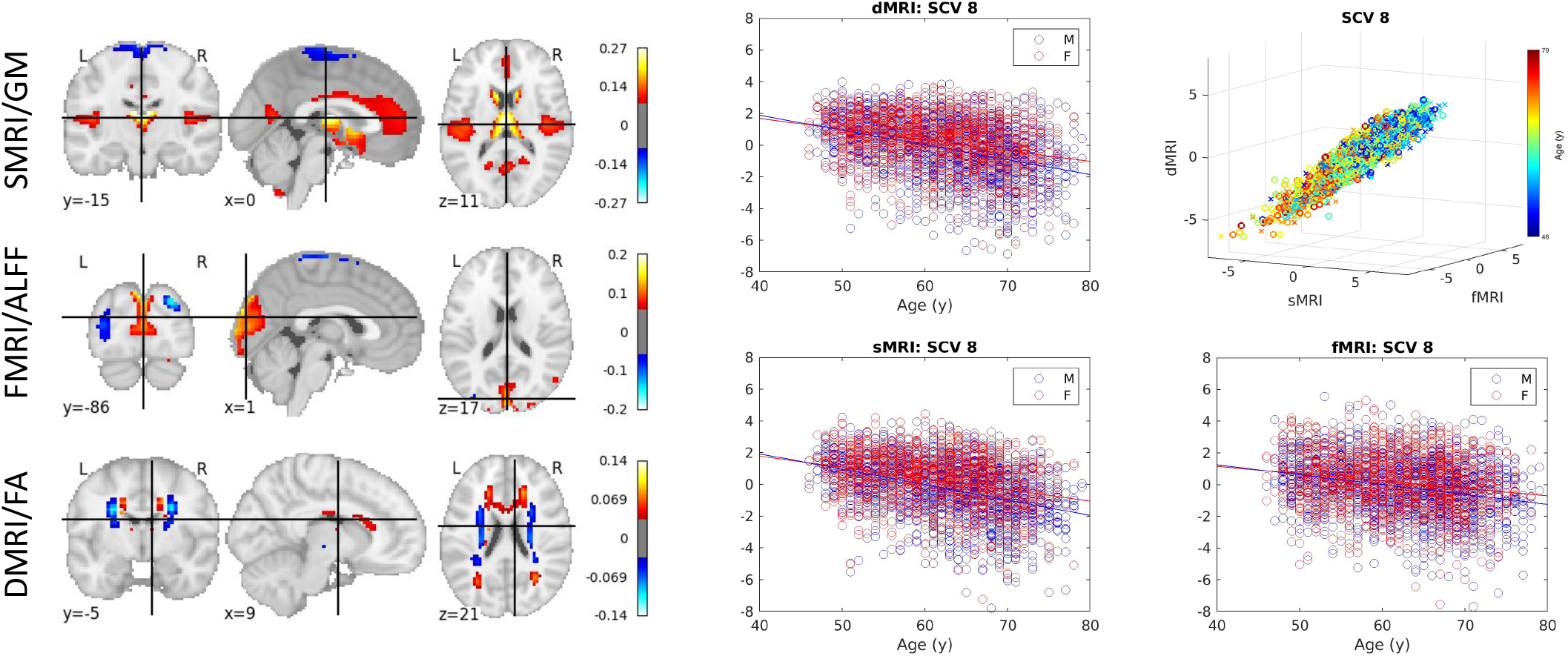
Age-related SCV. Left: The spatial maps corresponding to the mixing weights for each modality. Right: A 3D scatter plot of SCV 8 illustrates the strong association between multimodal subject expression levels, as well as their relationship to age, for the UK Biobank data. Each point represents a subject, color-coded by the age (circles indicate males; crosses indicate females). The same data as in the top right panel is depicted separately for each modality on the other panels, plotted against subject age.

The subject expression level sources from the SVD-shared information, sMRI and fMRI modalities in these five SCVs (15 sources in total) were subsequently used to predict the brain age (using chronological age as a surrogate) and estimate the brain-age delta. The mean absolute error of the overall brain-age delta estimation is 2.70 years when using the SVD-shared information from each SCV, plus the sMRI and fMRI sources (partialled over all fifteen sources) in one model. The mean absolute errors are 2.58, 0.41 and 0.61 years for models including only SVD-shared, partialled sMRI-only, partialled fMRI-only, respectively. Figure 4 (from top to bottom) shows 1) the predicted expression levels of each source as a function of a cubic age model (i.e., with linear, quadratic, and cubic age terms); 2) the correlation between age and the expression levels of SVD-shared features, sMRI and fMRI sources (and their partialled versions), as well as the regression coefficient and regression significance at the first stage; 3) the standard deviation of the estimated brain-age delta (and its partialled version) at the second stage, its mean absolute error, and the standard deviation of predicted age at the first stage. Ten out of the fifteen features have a *β*_1_ coefficient with significant age dependence (*p* < 0.05/15, Bonferroni correction). The bottom panel indicates that the SVD-shared features explain the largest amount of the age variance, with smaller but significant unique contributions from sMRI (SCVs 8, 16, 3, 17) and fMRI (SCV 5). In the same panel, the close agreement between standard deviations from predicted age and second stage brain-age delta suggests that the patterns extracted from the imaging data are largely descriptive of brain age, rather than chronological age.

**Figure 4:**
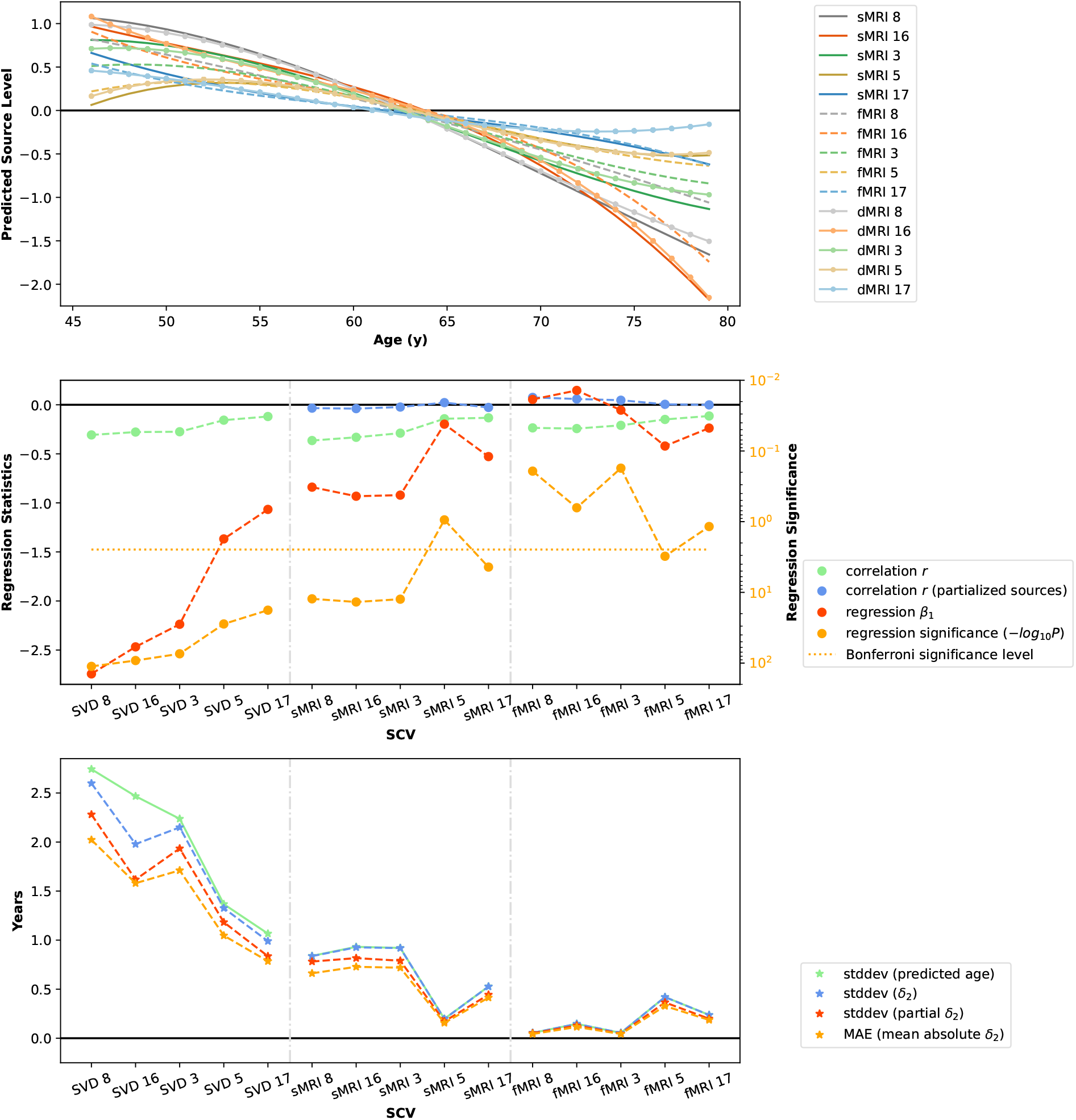
Brain-age delta modeling. Top: Predicted source levels as a function of chronological age using a cubic model. Middle: Regression statistics at stage 1 of brain-age delta modeling (see Equation 11). Bottom: Standard deviation of predicted age, standard deviation of stage 2 brain-age delta (see Equation 12) and its partialled version, and mean absolute error in stage 2 brain-age model. SCVs are sorted according to the correlation between their SVD-shared features and age. The close agreement between standard deviations from predicted age and second stage brain-age delta suggests that the patterns extracted from the imaging data are largely descriptive of brain age, rather than chronological age.

As presented in Figure 5, positive correlations between age delta and non-imaging variables indicate accelerated age decline. After FDR correction with q-value 0.05, a handful of statistically significant associations were identified between source-specific brain-age deltas and non-imaging phenotypes. Most associations were with time to answer in a prospective memory test, namely SCVs 5 (shared+sMRI), 17 (shared), and 8 (fMRI). Shared 5 and 17, as well as fMRI 8, indicate decelerated age decline with higher time to answer. sMRI 5 indicates accelerated age decline with higher time to answer. Only shared 5 and 17 associations seem to replicate significantly within males and females. Also, higher frequency of exercise in last 4 weeks associated with accelerated age decline in fMRI 17.

**Figure 5:**
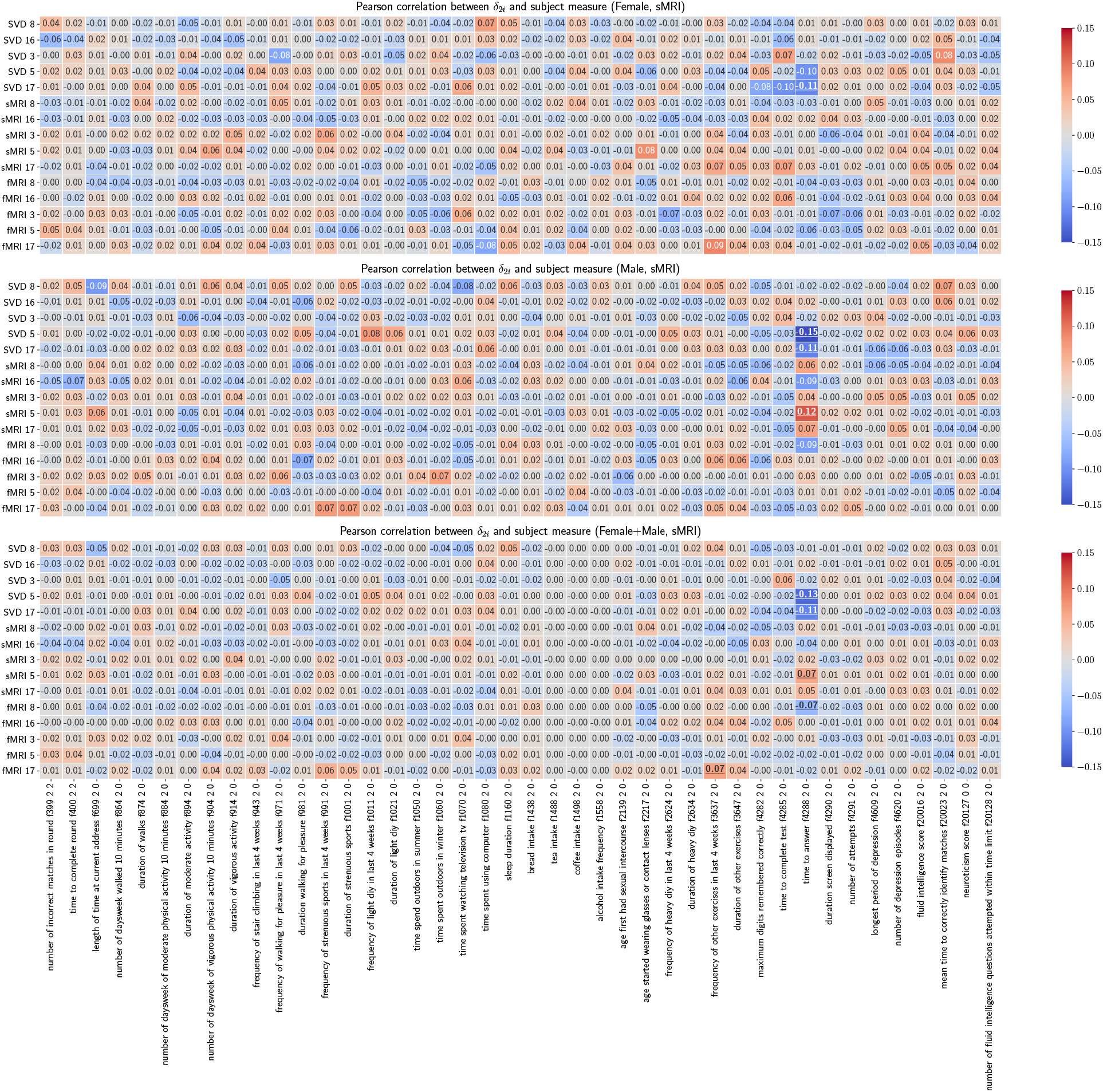
Pearson correlations between *δ*_2*i*_ vectors and subject measures. Pearson correlations are calculated between 15 *δ*_2*i*_ vectors and 42 variables for each gender and for all subjects. The underscored correlations are significant.

### 3.2. Associations with schizophrenia

Linked subject expression levels were also identified in the patient dataset. In particular, SCV 19 (see Figure 6) not only shows association with age, but also provides evidence of age-adjusted group differences between controls and patients with schizophrenia. Specifically, patients show lower source levels in all three modalities. Moreover, SCV 19 remarkably replicates the spatial patterns of UKB SCV8 described in Section 3.1.

**Figure 6:**
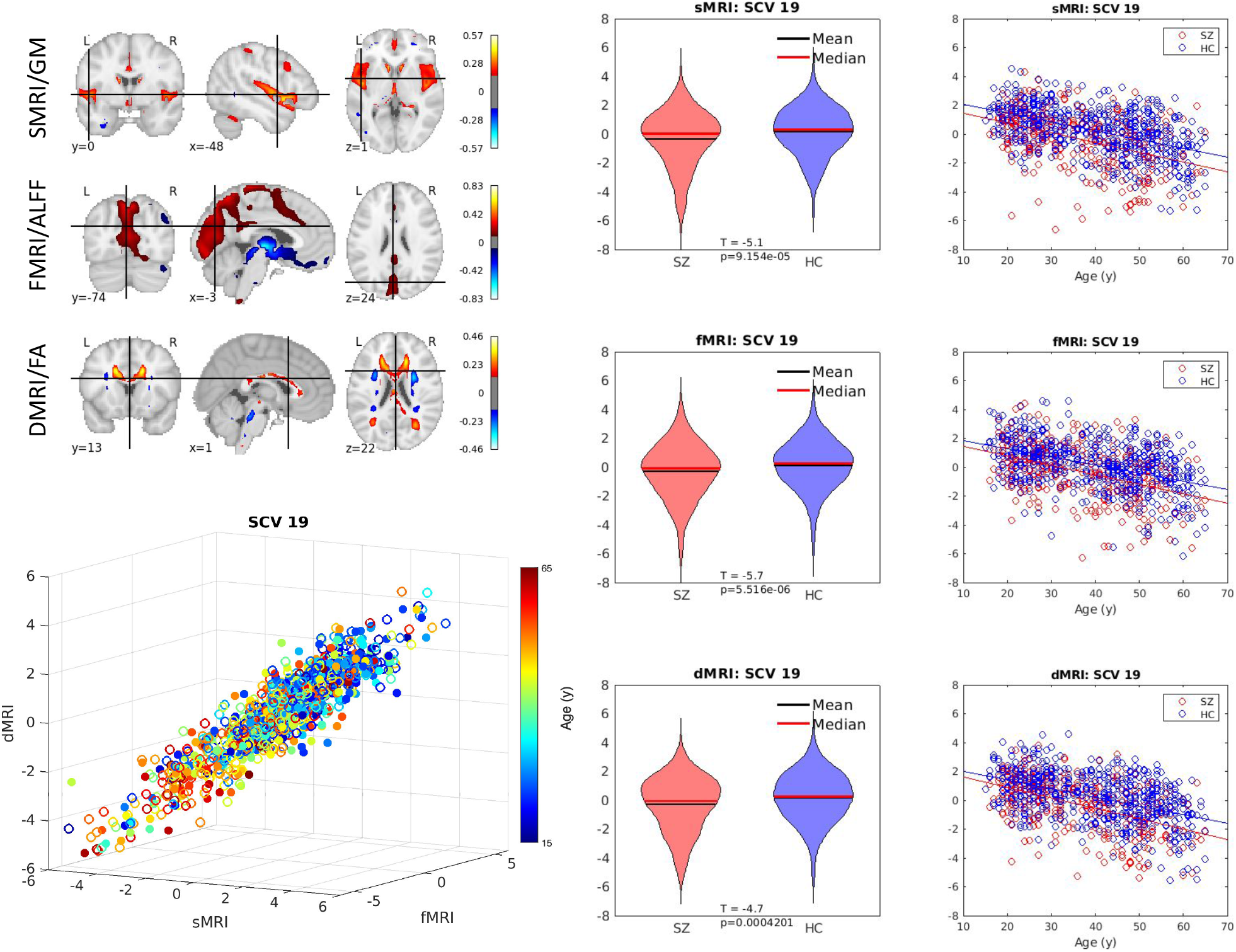
Schizophrenia-related SCV. Top left: The spatial maps correspond to the mixing weights for each modality. Bottom left: A 3D scatter plot of SCV 19 illustrates the association between the identified multimodal subject expression levels and age for patient data color coded by subject age (circles indicate females; filled circles indicate males). Right: The SCVs are also plotted by control (HC) and schizophrenia patient group (SZ) as violin plots and by age as scatter plots to demonstrate consistent reduction in source intensities in SZ patients across all ages.

In addition, SCV 14 (see supplementary section Figure 15) is also identified as a schizophrenia-related source. In SCV 14, patients have higher source intensities in all three modalities. In sMRI, negative weights can be found in the insula and the frontal inferior operculum while positive weights are shown in the temporal lobe, the occipital lobe, the lingual gyrus and the orbital part of inferior frontal gyrus. In fMRI, the superior frontal cortex and the parietal superior cortex have positive activation. In dMRI, positive weights can be seen in the superior longitudinal fasciculus while negative weights can be found in the cingulum. Positive weights indicate accelerated brain aging.

### 3.3. Associations with sex effects in UK Biobank

Five SCVs (9, 10, 16, 17 and 22) show significant sex effects in UK Biobank data. Females show significantly higher source intensities than males in SCV 9 and 16 (see Figure 7 for SCV 16) while males show significantly higher source intensities than females in SCVs 10, 17 and 22 (see supplementary section, Figure 14).

**Figure 7:**
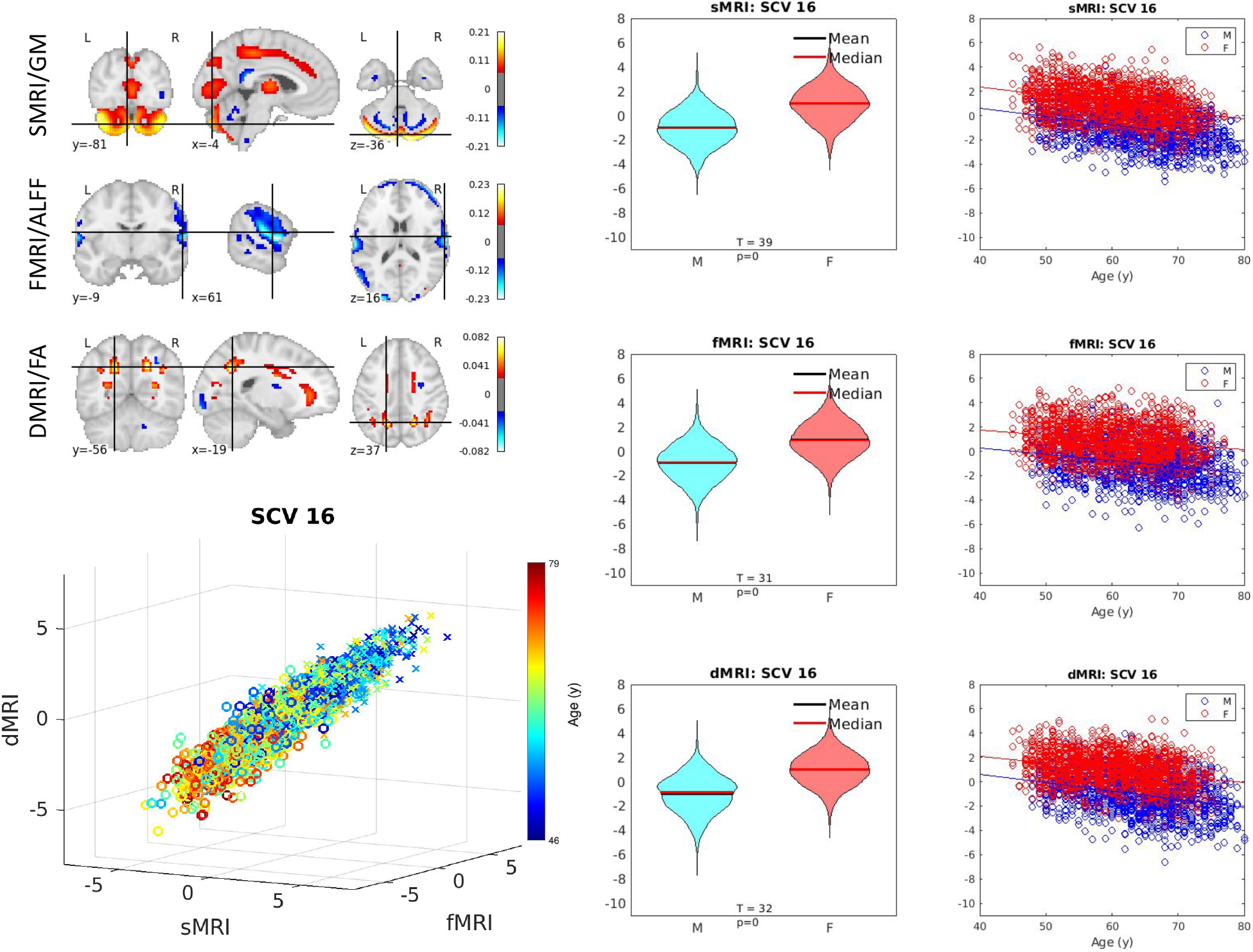
Sex-related SCV. Illustration of SCV 16. Top left: The spatial maps correspond to the mixing weights for each modality. Bottom left: Multimodal subject expression level sources, depicting all subjects (colored by age; circles indicate females while crosses indicate males). Right: Source intensity differences related to sex effects.

For SCV 16, in both sexes, notice the linear covariation with age and similar trajectory of decline with age observed in cerebellar regions (sMRI/GM), parietal cortex (fMRI/ALFF), and parietal cortico-pontine tracts (dMRI/FA).

In sMRI, SCVs 10 and 17 show negative weights in the parietal lobe and the cerebellum, respectively. SCV 9 shows positive weights in the temporal lobe and negative weights in the angular gyrus. SCV 22 shows both positive and negative weights in the cerebellum. In fMRI, SCV 16 shows negative weights in the temporal lobe, SCVs 10 and 17 show positive weights in the frontal lobe. SCV 9 has positive weights in the cuneus in the occipital lobe. SCV 22 shows positive weights in the olfactory cortex. In dMRI, SCVs 10 and 16 have positive weights in caudate nucleus, while SCV 17 shows negative weights in the posterior cingulate cortex. SCV 9 and SCV 22 show positive weights in the vertical occipital fasciculus and the brain stem, respectively. As the age increases, regions with positive weights experience decline (conversely, increase in regions with negative weights).

### 3.4. Associations with cognitive performance in UK Biobank

As shown in Figure 8, the SCV 22 shows significant linear association with time to answer (TTA) in a prospective memory test across all three modalities. Subjects with faster responses have higher source intensities. Males have slightly higher source intensities than females in all three modalities. The spatial maps highlight the cerebellum in sMRI, the olfactory cortex in fMRI, and the brain stem in dMRI.

**Figure 8:**
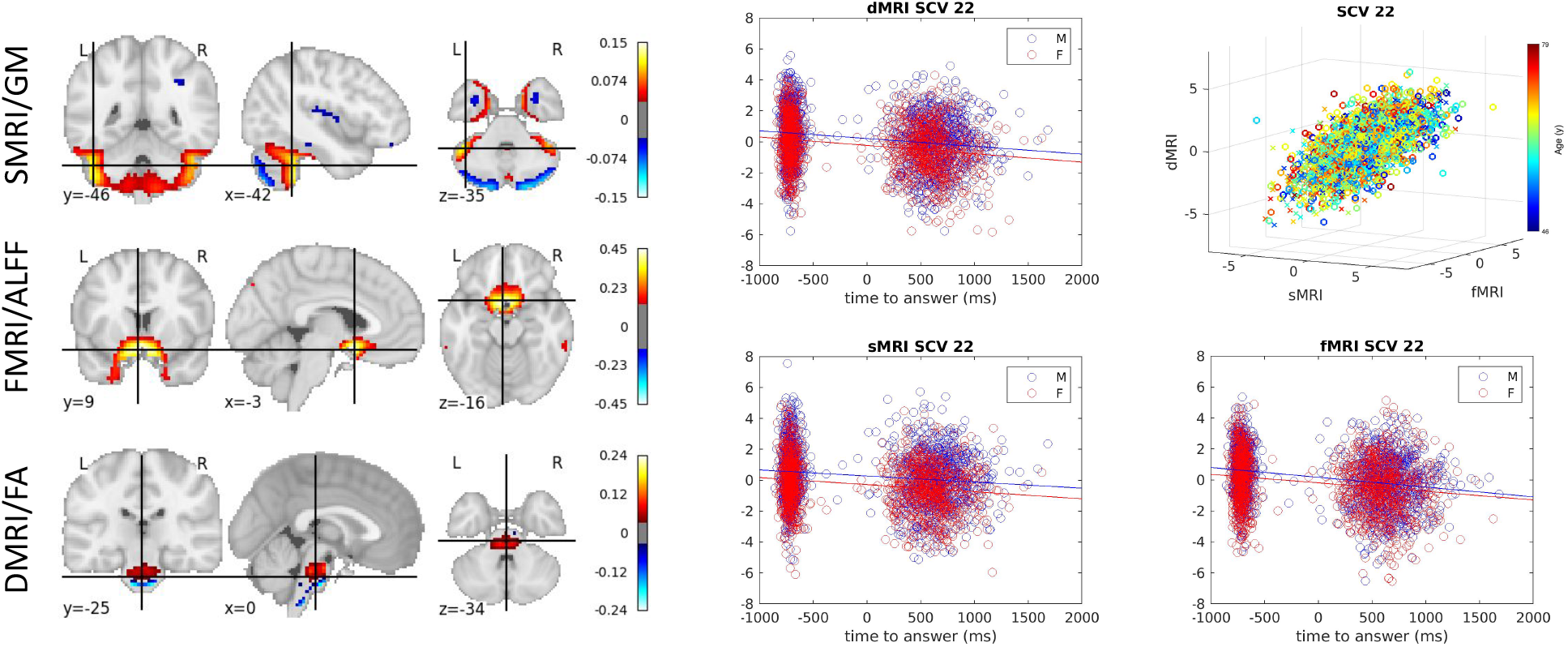
Time to answer-related SCV. Left: The spatial maps corresponding to the mixing weights for each modality are shown. Right: A 3D scatter plot of SCV 22 illustrates the association between the identified multimodal subject expression levels and age for UK Biobank data. 2D scatter plots illustrate the relationship between the time to answer and the source intensities for each of the three modalities.

## 4. Discussion

In this work, we demonstrate that the proposed multimodal IVA model, initialized with a multimodal group projection (see Methods 2.4), can extract independent multimodal subspaces that show significant covariation with subject phenotypes. The proposed approach directly leverages multimodal associations and is applied to two large datasets independently. The results highlight several linked multimodal patterns of expression over subjects that are significantly associated with aging, sex, cognition, and psychosis indicators.

Five SCVs (SCVs 3, 5, 8, 16, 17) were identified that were significantly associated with age based on effect size (*ϵ*^2^). As presented in Figure 3, for UK Biobank dataset, age associated decline in gray matter density (hot spatial areas of sMRI SCV 8 weights) was primarily seen in caudate, thalamus, insular regions, anterior and posterior cingulate cortex, and lingual gyrus, consistent with earlier findings (Brickman et al., 2007). ALFF maps corresponding to SCV 8 suggest reductions in parietal and visual regions of the brain that covary with the structural changes. In dMRI, the identified spatial map weights show FA intensity reductions in periventricular regions, including superior and posterior thalamic radiation, are associated with the age.

Brain-age delta was estimated using 5 shared-SVD source features, 5 partialled sMRI sources and 5 partialled fMRI sources. Each individual source from each of the three modalities within an SCV describes a pattern of subject-specific expression levels that is linked (covaries) between modalities within that SCV, while being statistically independent from sources in other SCVs. The mean absolute error of overall age prediction is 2.70 years, and the mean absolute errors are 2.58, 0.41, and 0.61 years when using only the 5 individual sources from each feature type (shared-SVD, partialled sMRI, and partialled fMRI, respectively) — both cases based on the same multimodal IVA decomposition. The association between each individual source and the age for each subject was also computed. As shown in Figure 4, all five SVD-shared features, four sMRI sources (SCVs 3, 8, 16, 17), and one fMRI source (SCV 5) show significant age regression beta weights. Though the unique variability of the partialled sMRI source from SCV 5 does not contribute significantly to the age regression model, the corresponding partialled fMRI source from SCV 5 does show significant contribution. Thus, all five SCVs display significant age regression coefficients in at least one modality, and we conclude that each modality contributes complementary information not captured in other modalities. Moreover, the brain-age delta result using only the 5 sources from shared-SVD features supports that each modality is valuable and largely predictive of brain age, which is largely a consequence of the inherent multimodal link (correlation) among expression levels within SCVs. The advantage of such multimodal analysis is not only that it leverages hidden covariation across modalities to recover linked subject expression patterns, but also that it captures unique information from each modality by allowing these patterns to differ across modalities. Note that in our brain-age delta analysis, confound variables head size, scanner table position and scan-date-related slow drifts described in Smith et al. (2020) were not included.

Two SCVs (SCVs 14, 19) in the patient dataset are associated with schizophrenia effects. SCV 19 revealed schizophrenia effects as sMRI gray matter intensity reduction in the temporal lobe, consistent with previous findings (Karlsgodt et al., 2010). Schizophrenia-associated reductions can also be observed in the medial parietal lobe, including the posterior cingulate cortex in fMRI ALFF maps, also aligned with previous literature (Venkataraman et al., 2012). The dMRI spatial map shows decreased FA values in the anterior thalamic radiation and forceps minor tracts, which is close to the superior longitudinal fasciculus and cingulate bundle reported in Kyriakopoulos et al. (2008). As is the nature of the proposed multimodal IVA approach, all of these regional changes covary at similar levels over subjects across all three modalities.

Sex differences were mainly reflected in SCVs 9, 10, 16, 17 and 22. Females show significantly higher source intensities than males in SCV 16 while males show higher intensities than females in SCV 10 and 17. Consistent with previous studies (Liu et al., 2020; Ritchie et al., 2018), males have significantly higher source intensities than females in the parietal lobe (SCV 10) and the occipital lobe (SCV 17) from sMRI, the frontal lobe in both SCV 10 and 17 from fMRI, the corpus callosum and the cingulum in both SCV 10 and SCV 17 from dMRI. As shown in Figure 7 for SCV 16, females have higher intensities than males in the occipital lobe from sMRI, the temporal lobe from fMRI and the cingulum from dMRI. The decline with aging association can be found in the sMRI and dMRI modalities while the opposite association can be observed in the temporal lobe from fMRI ALFF maps.

One SCV (SCV 22) has associations with time to answer in a prospective memory test. As presented in Figure 8, the cerebellum identified by the sMRI source has been found to relate to the biological basis of time perception and fast response (Grondin, 2010), and prospective memory (Cona et al., 2016).

In summary, we demonstrate the ability of multimodal independent vector analysis to extract linked multimodal modes of subject variations that capture different aspects of phenotypical information including aging effects, schizophrenia-related biomarkers, sex effects and cognitive performance across two large independent datasets. With the increasing demand of multimodal neuroimaging data analysis, the MMIVA fusion model can be applied to identify linked sources with associated phenotypes across multiple neuroimaging modalities and multiple datasets.

## 5. Acknowledgments

This work was supported by NIH grant R01MH118695 and NSF grant #2112455.

Dr. Ford is the recipient of a Distinguished Research Career Scientist award (IK6CX002519) from the Department of Veterans Affairs.

This research has been conducted using the UK Biobank Resource under Application Number 34175.

## 6. Supplementary Materials

### 6.1. Phenotype Variables

54 phenotype variables were further reduced to 28 variables.

28 physical exercise variables were decomposed to 8 principal components by PCA. 2 age-related variables (“years since first sexual intercourse 2 0” and “years since started wearing glasses 2 0”) were dropped because they are highly correlated with other age variables. 2 variables associated with fluid intelligence (“number of fluid intelligence questions attempted within time limit” and “interaction between number of attempts and score”) and 4 other log variables (time to complete round f400 2 2, log time to answer, inverse log duration screen displayed, inverse log number of attempts) were also removed, resulting in 26 variables in total.

The 26 variables are common to all of the 3 modality-specific MANCOVA models. The interaction between age and sex is also included for each of 3 modality-specific MANCOVA models. In addition to the 27 common variables, sMRI and dMRI modalities have 1 nuisance variable (rspatialNorm: correlation between individual GM/FA map to the site-specific mean) and fMRI has 2 nuisance variables (rspatialNorm, and meanFD: estimated mean framewise displacement of the subject during the scan).

**Table 3:** UK Biobank Phenotype Variables.

| Variable ID | Variable Name |
| --- | --- |
| f399 2 2 | number of incorrect matches in round |
| f400 2 2 | time to complete round |
| f699 2 0 | length of time at current address |
| f864 2 0 | number of daysweek walked 10 minutes |
| f874 2 0 | duration of walks |
| f88 | number of daysweek of moderate physical activity 10 minutes |
| f894 2 0 | duration of moderate activity |
| f90 | number of daysweek of vigorous physical activity 10 minutes |
| f914 2 0 | duration of vigorous activity |
| f943 2 0 | frequency of stair climbing in last 4 weeks |
| f971 2 0 | frequency of walking for pleasure in last 4 weeks |
| f981 2 0 | duration walking for pleasure |
| f991 2 0 | frequency of strenuous sports in last 4 weeks |
| f1001 2 0 | duration of strenuous sports |
| f1011 2 0 | frequency of light diy in last 4 weeks |
| f1021 2 0 | duration of light diy |
| f1050 2 0 | time spend outdoors in summer |
| f1060 2 0 | time spent outdoors in winter |
| f1070 2 0 | time spent watching television tv |
| f1080 2 0 | time spent using computer |
| f1160 2 0 | sleep duration |
| f1438 2 0 | bread intake |
| f1488 2 0 | tea intake |
| f1498 2 0 | coffee intake |
| f1558 2 0 | alcohol intake frequency |
| f2139 2 0 | age first had sexual intercourse |
| f2217 2 0 | age started wearing glasses or contact lenses |
| f2624 2 0 | frequency of heavy diy in last 4 weeks |
| f2634 2 0 | duration of heavy diy |
| f3637 2 0 | frequency of other exercises in last 4 weeks |
| f3647 2 0 | duration of other exercises |
| f4288 2 0 | time to answer |
| f4609 2 0 | longest period of depression |
| f20016 2 0 | fluid intelligence score |
| f20023 2 0 | mean time to correctly identify matches |
|  | number of fluid intelligence questions attempted within time |
| f21003 2 0 | age when attended assessment centre |
| 2 0 | total hours walked 10 minutes |
| 2 0 | total hours moderate physical activity 10 minutes |
| 2 0 | total hours vigorous physical activity 10 minutes |
| 2 0 | total hours of walking for pleasure in last 4 weeks |
| 2 0 | total hours of strenuous sports in last 4 weeks |
| 2 0 | total hours of other exercises in last 4 weeks |
| 2 0 | total hours of light diy in last 4 weeks |
| 2 0 | total hours of heavy diy in last 4 weeks |
| 2 0 | number of physical activities wrt walking for pleasure |
| 2 0 | years since first sexual intercourse |
| 2 0 | years since started wearing glasses |
| 2 0 | log time to answer |
| 2 0 | inverse log duration screen displayed |
| 2 0 | inverse log number of attempts |
| 2 0 | log pm score |
| 2 0 | fluid intelligence interaction |
| f31 0 0 | sex |
| f22001 0 0 | genetic sex |

### 6.2. Aging Effects

Five SCVs (3, 5, 8, 16, 17) were significantly associated with age identified by the MANCOVA analysis.

**Figure 9:**
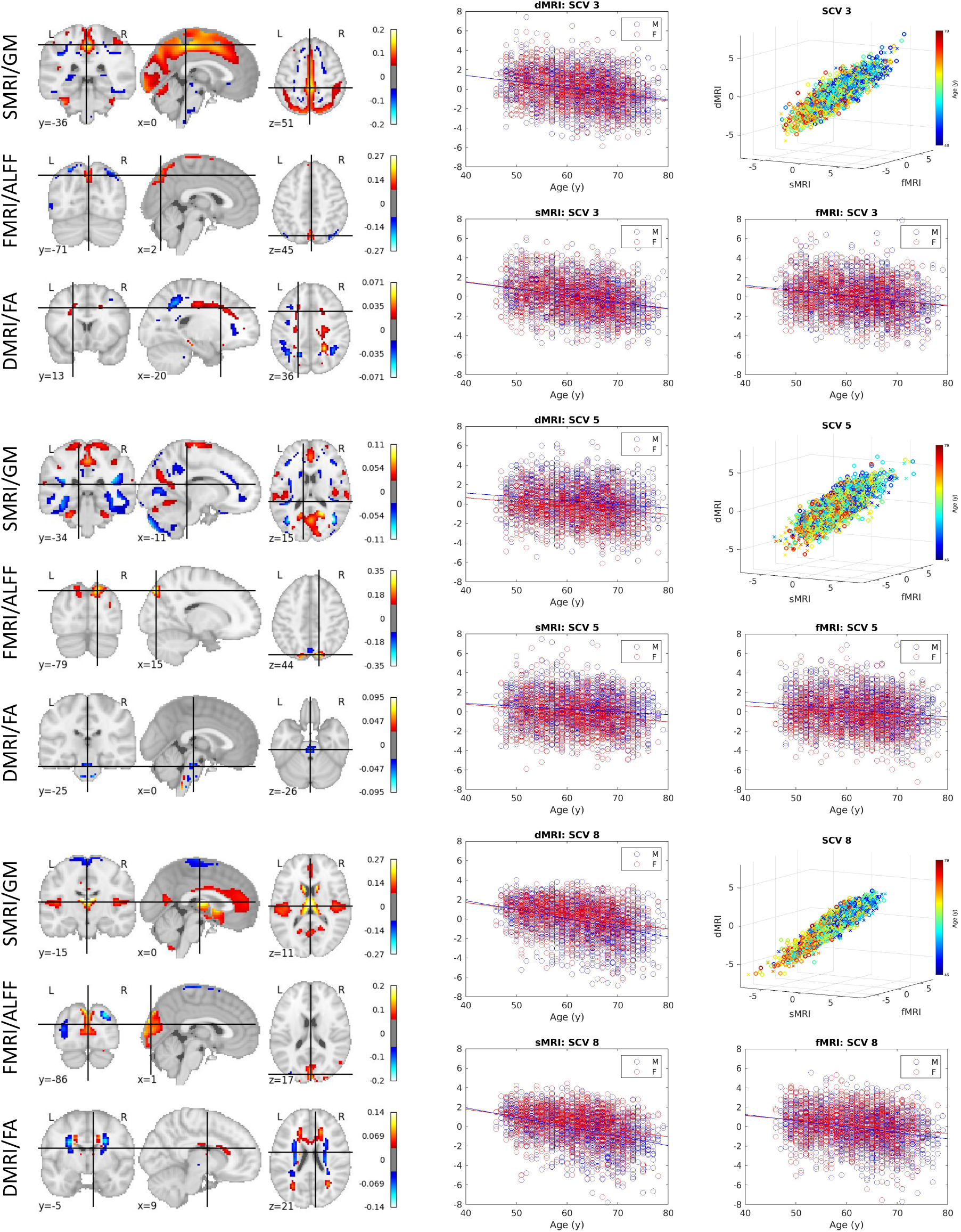
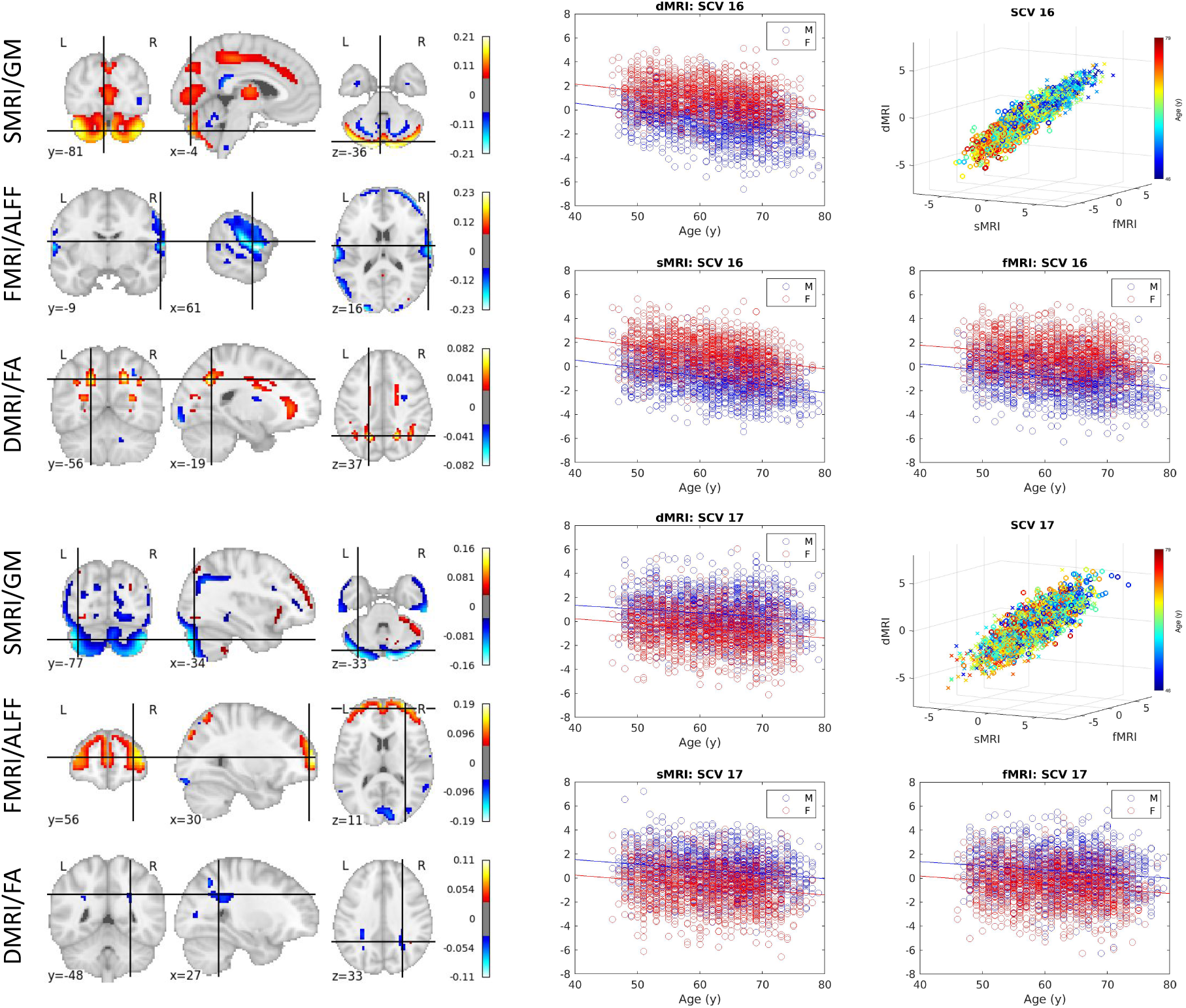
Age-related SCV. Right: 3D scatter plots of SCVs 3, 5, 8, 16, 17 illustrate the association between the identified multimodal subject expression levels and age for UK Biobank data. Each point represents a subject, color coded by the age (circles indicate females; crosses indicate males). Left: The spatial maps corresponding to the mixing weights for each modality are also shown.

### 6.3. Brain-age Delta Modeling

**Figure 10:**
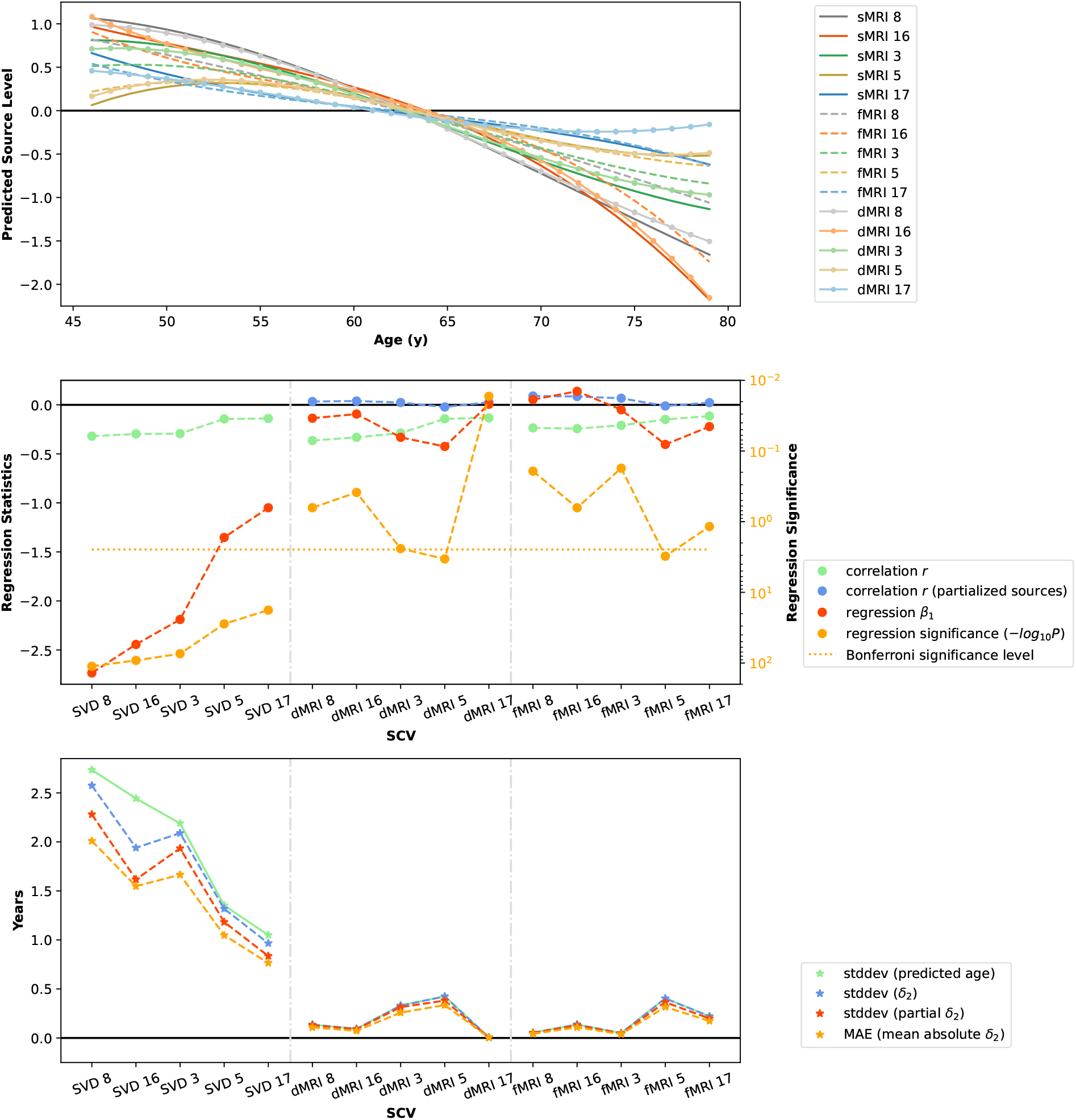
Brain age delta modeling (dMRI). Top: Predicted source levels as a function of brain age using a cubic model. Middle: Regression statistics at stage 1 of brain-age delta modeling (*age* = *S* * ***β***_1_). Bottom: Standard deviation of predicted age, standard deviation of estimated stage 2 brain-age delta, and mean absolute error in stage 2 brain-age model. SCVs are sorted according to the correlation between the age and the source within each modality.

**Figure 11:**
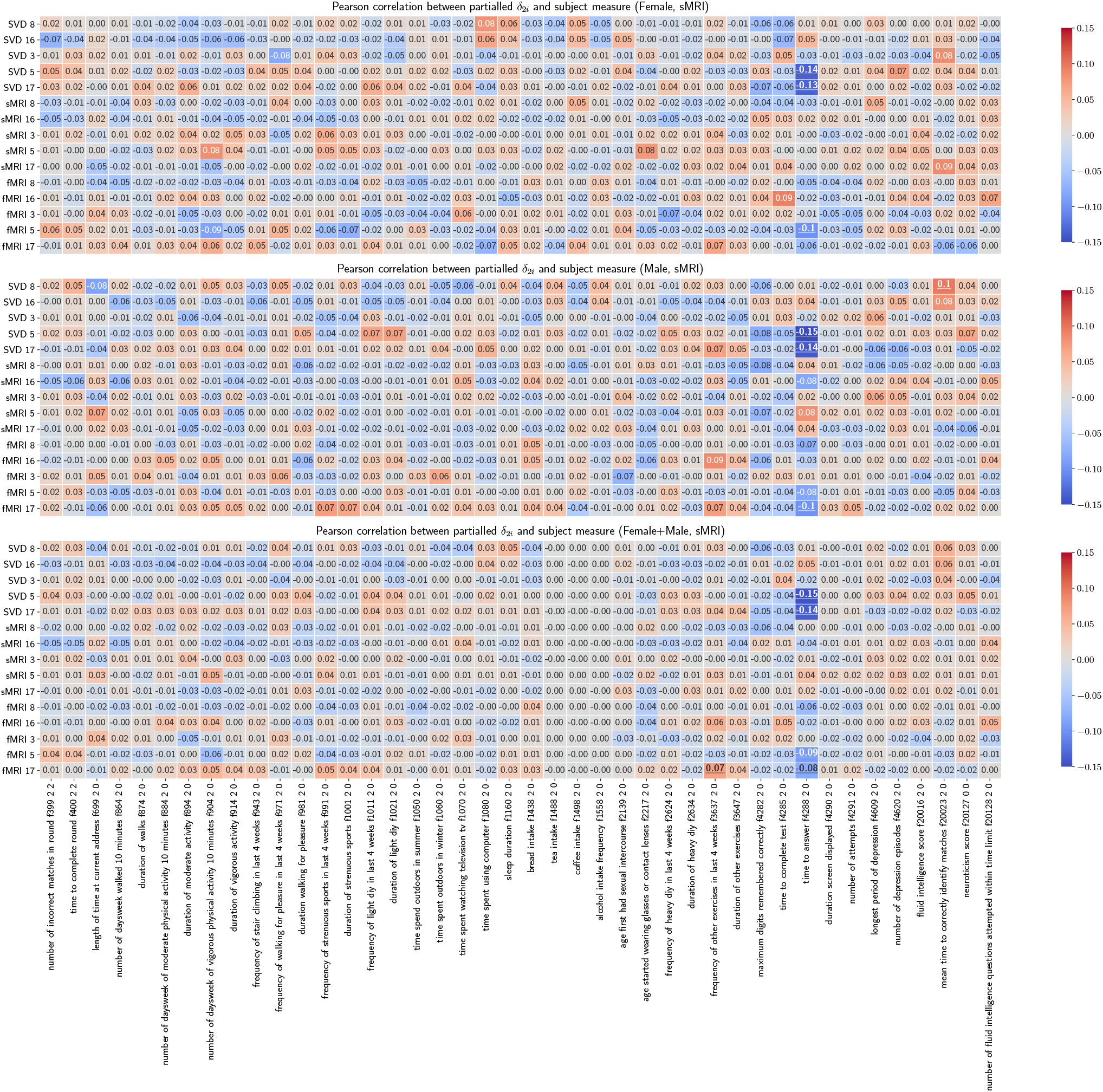
Pearson correlations between partialled *δ*_2*i*_ vectors (sMRI) and subject measures. Pearson correlations are calculated between 15 partialled *δ*_2*i*_ vectors and 42 variables for each gender and for all subjects. The underscored correlations are significant.

**Figure 12:**
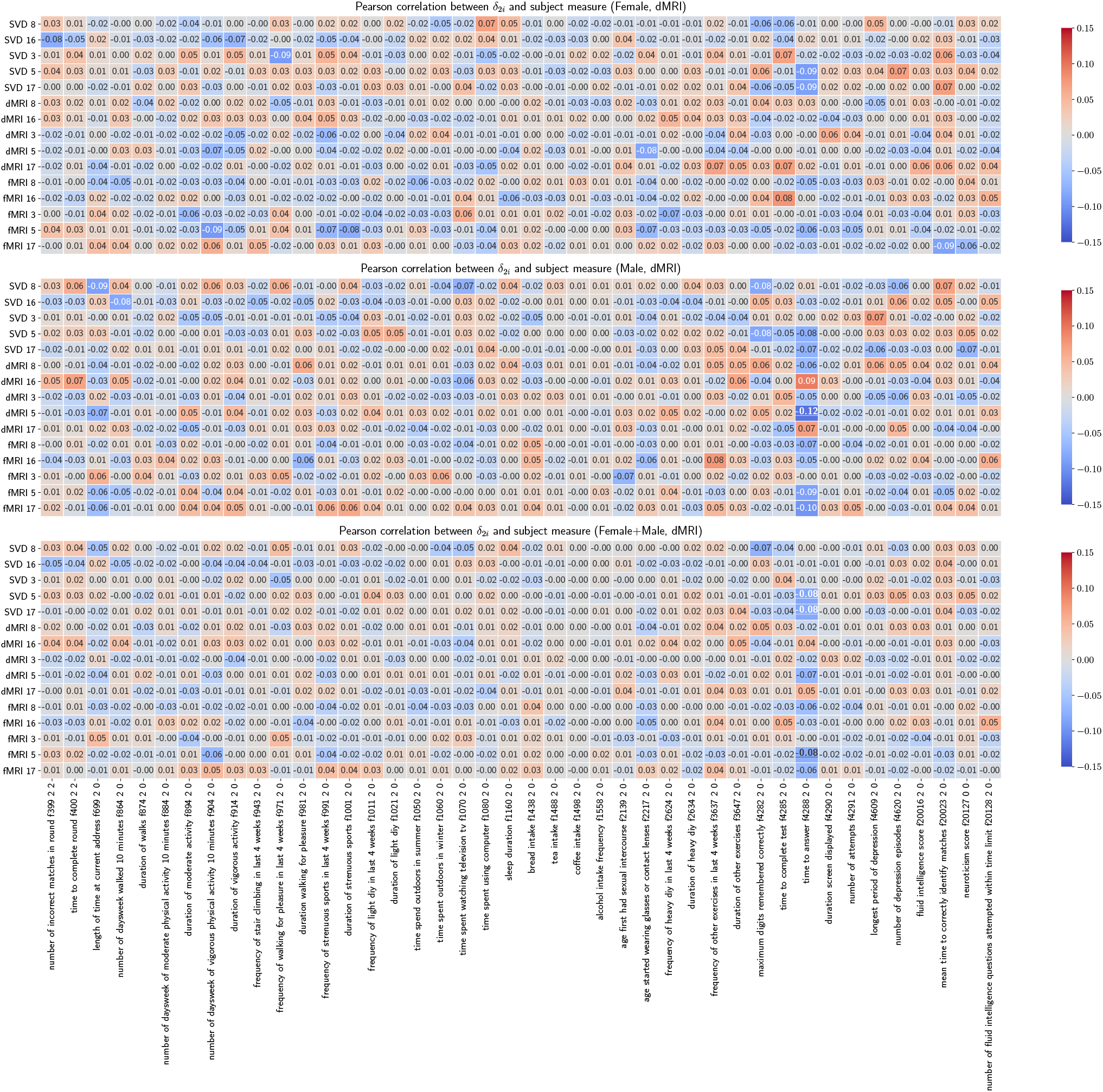
Pearson correlations between *δ*_2*i*_ vectors (dMRI) and subject measures. Pearson correlations are calculated between 15 *δ*_2*i*_ vectors and 42 variables for each gender and for all subjects. The underscored correlations are significant.

**Figure 13:**
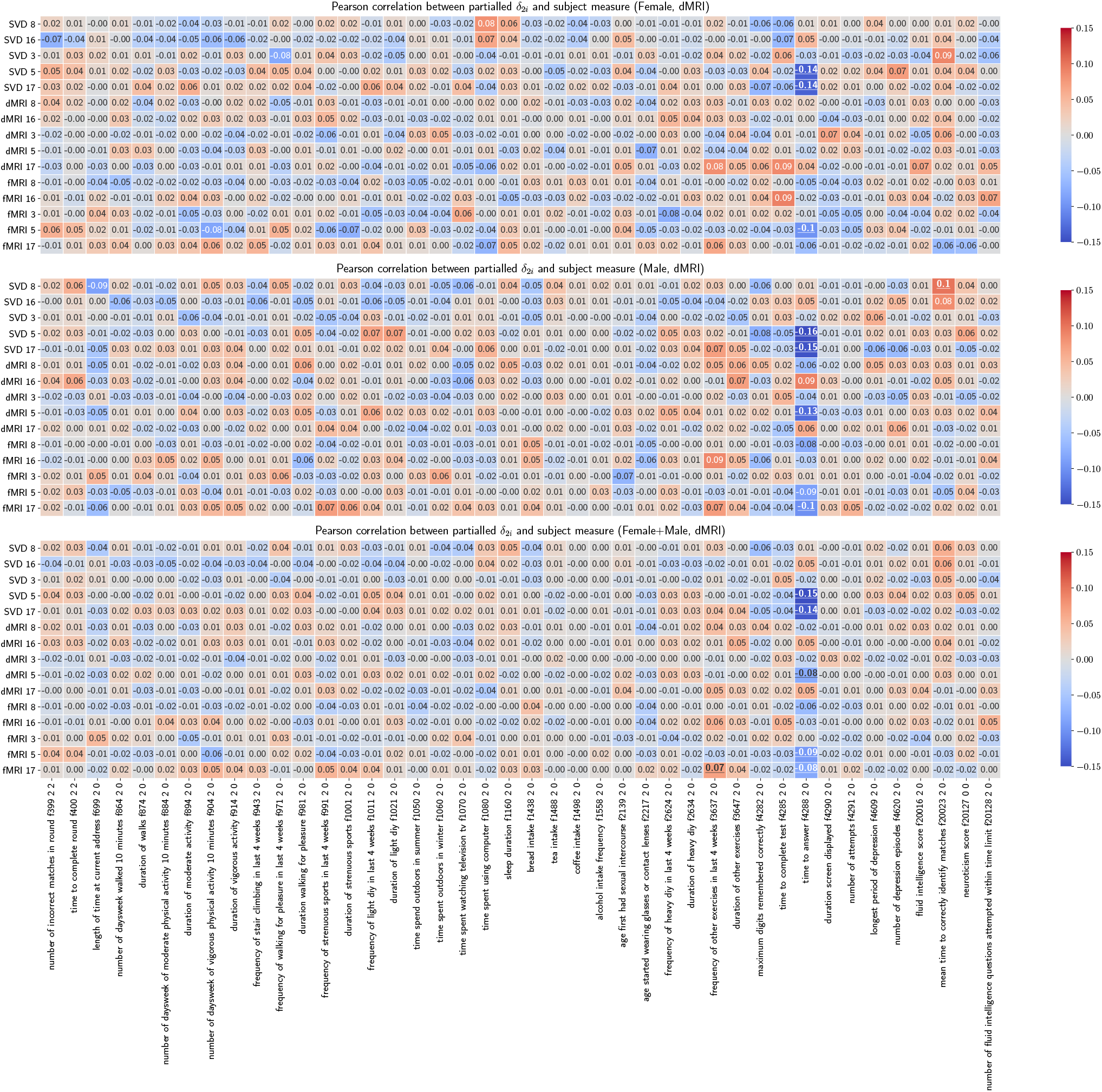
Pearson correlations between partialled *δ*_2*i*_ vectors (dMRI) and subject measures. Pearson correlations are calculated between 15 partialled *δ*_2*i*_ vectors and 42 variables for each gender and for all subjects. The underscored correlations are significant.

### 6.4. Sex Effects

SCVs (9, 10, 17, 22) were significantly associated with sex identified by the MANCOVA analysis and the effect size measure.

**Figure 14:**
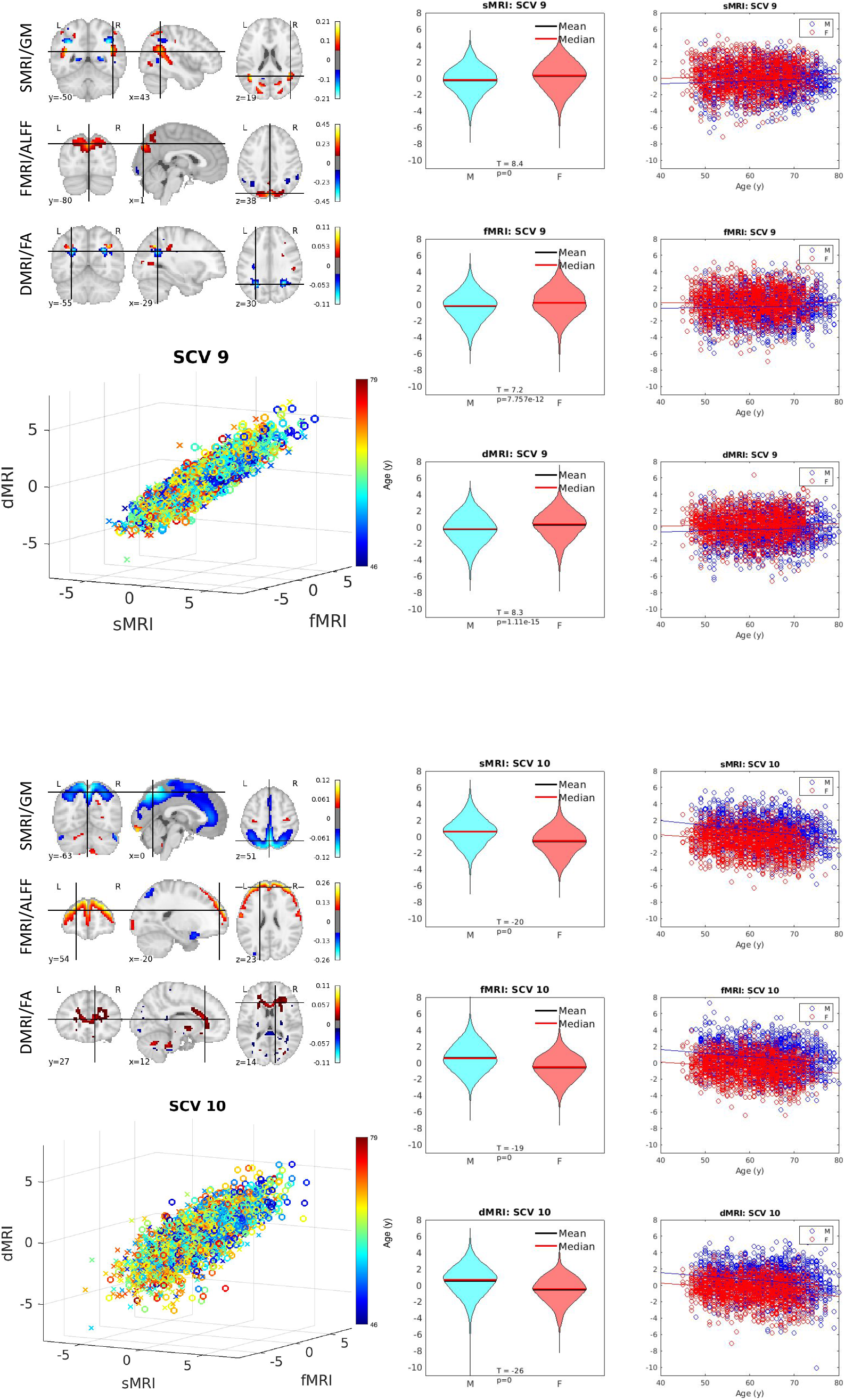
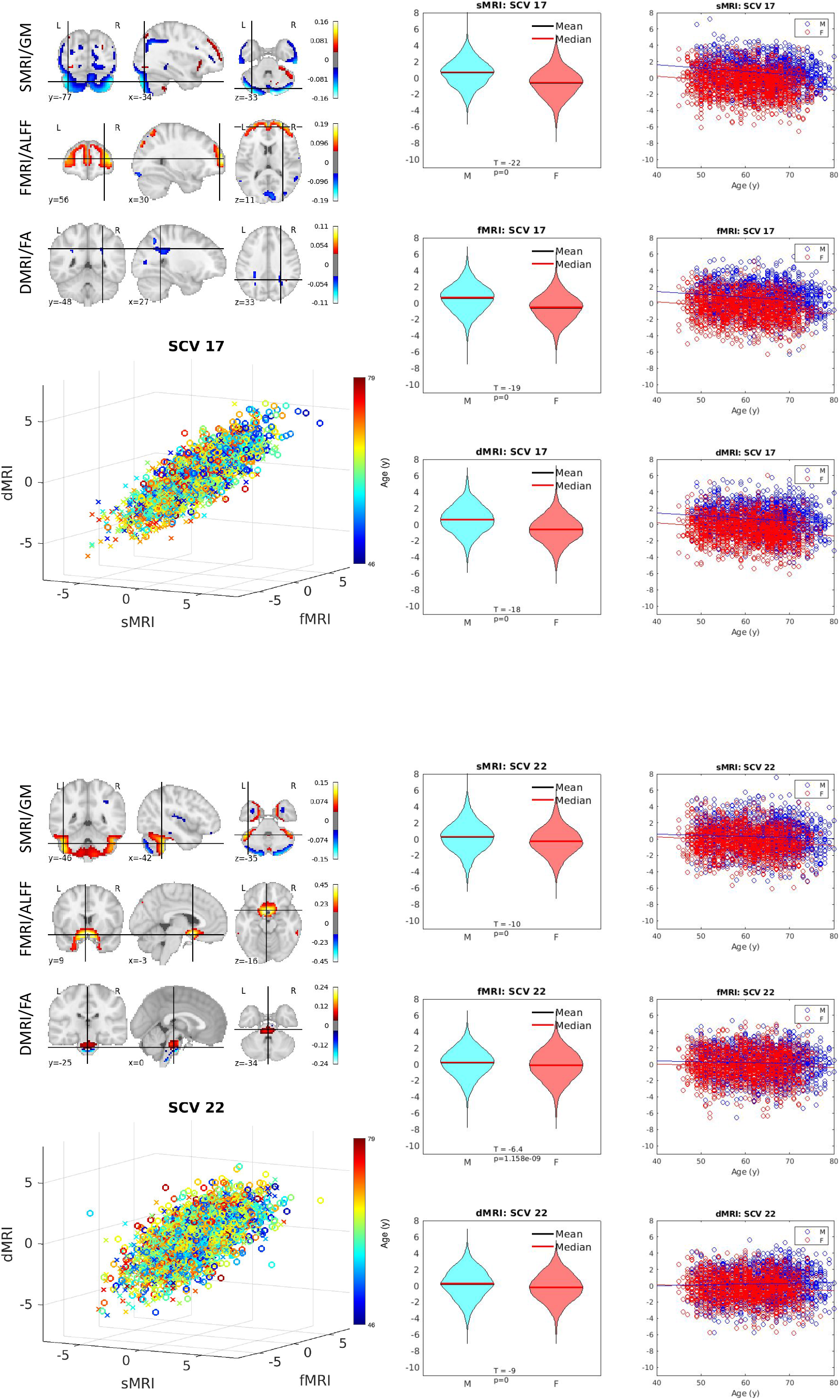
Sex-related SCV. Illustration of SCVs 9, 10, 17 and 22. Top left: Spatial maps. Bottom left: Multimodal subject expression level sources, depicting all subjects (colored by age). Right: Source intensity differences related to sex effects.

### 6.5. Schizophrenia Effects

**Figure 15:**
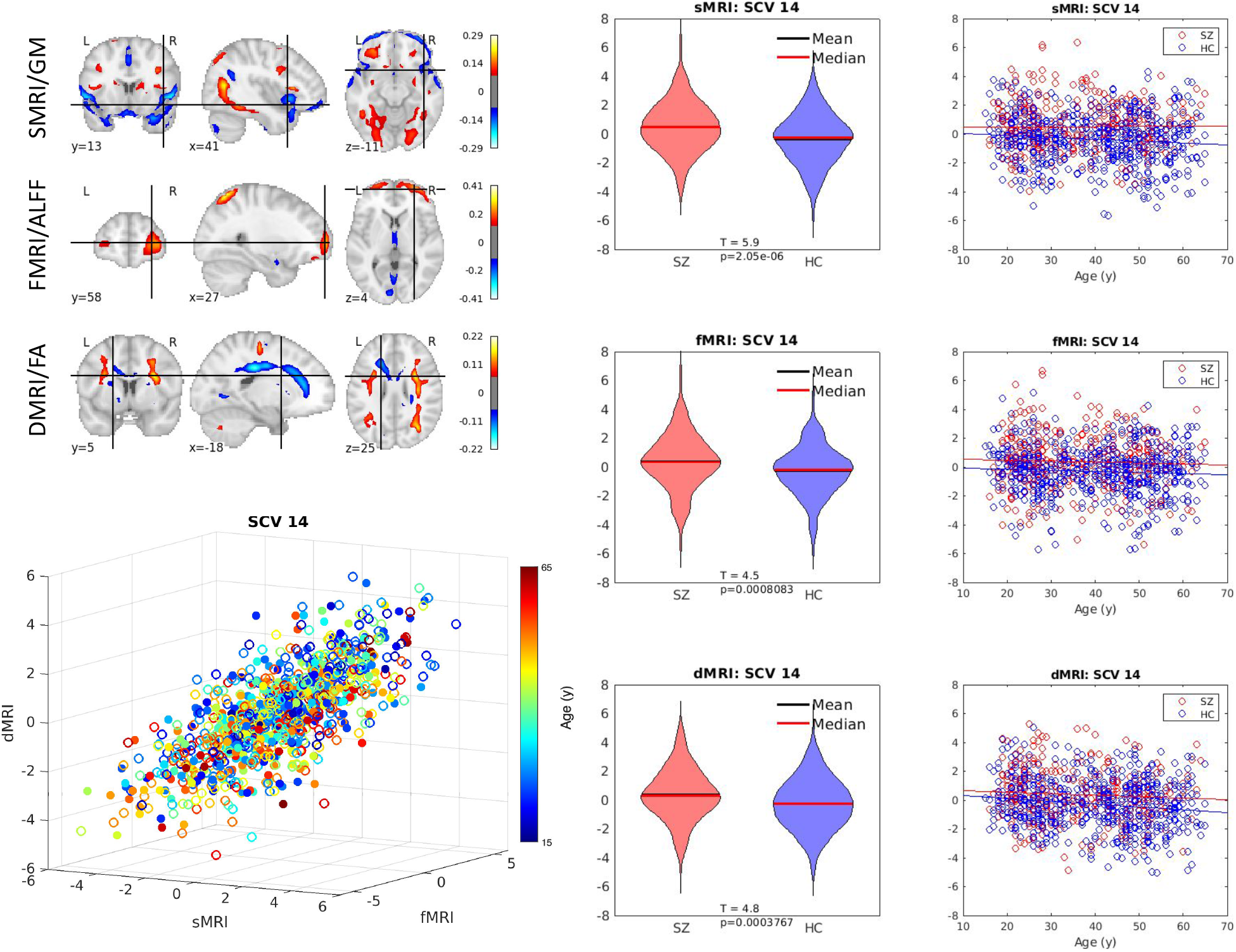
Schizophrenia-related SCV. Illustration of SCV 14. Top left: Spatial maps. Bottom left: Multimodal subject expression level sources, depicting all subjects (colored by age). Right: Source intensity differences related to schizophrenia effects.

### 6.6. Effect Size Measure (UK Biobank Dataset)

SCVs that have significant interactions and are replaced by Type III measure are listed below:

sMRI: 6, 11, 18, 21, 23, 24, 28

fMRI: 18, 25

dMRI: 6, 18, 22, 26

Red star is for p-values, blue dot is for effect size. Horizontal lines indicate effect size threshold for small, medium, and large.

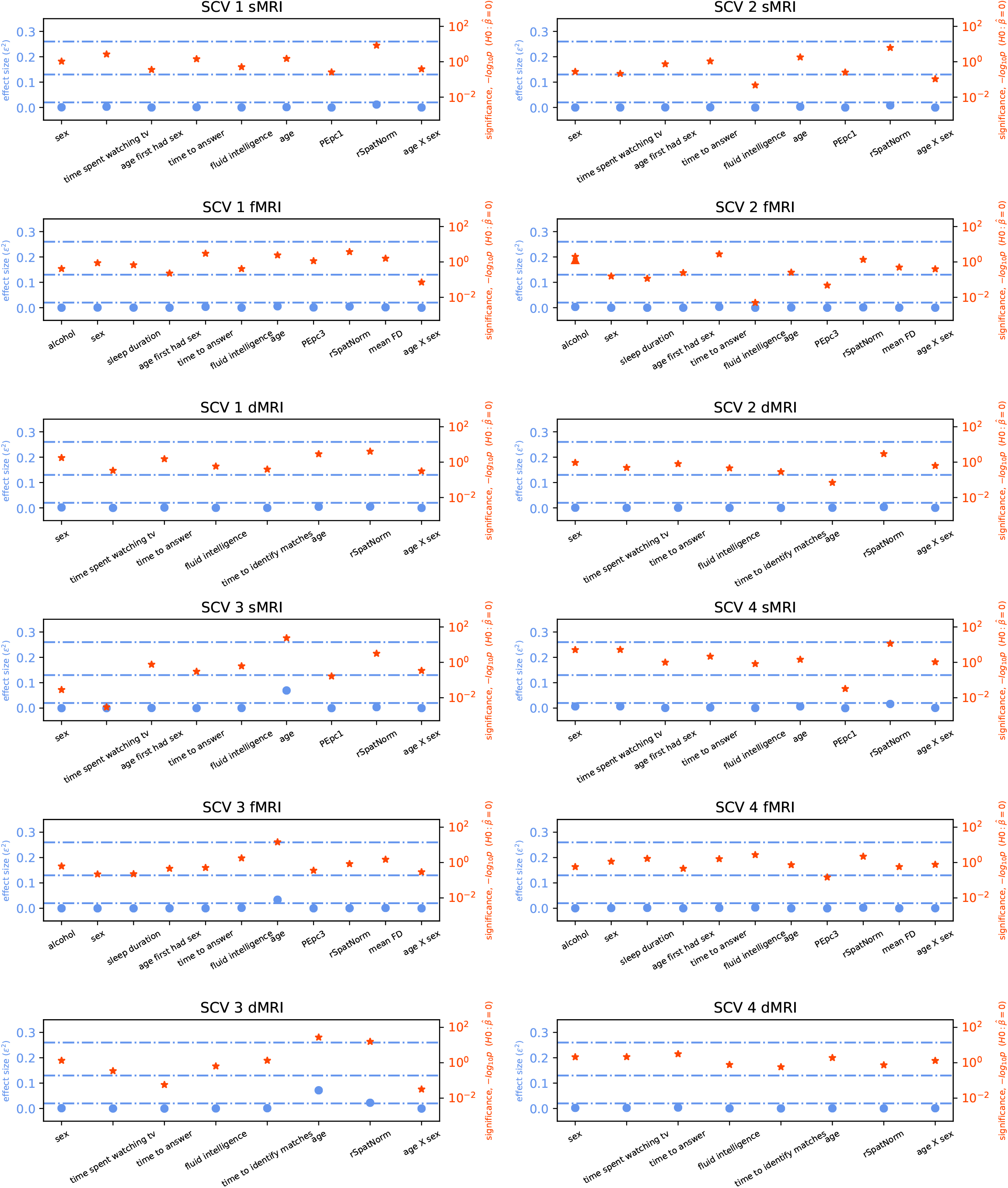

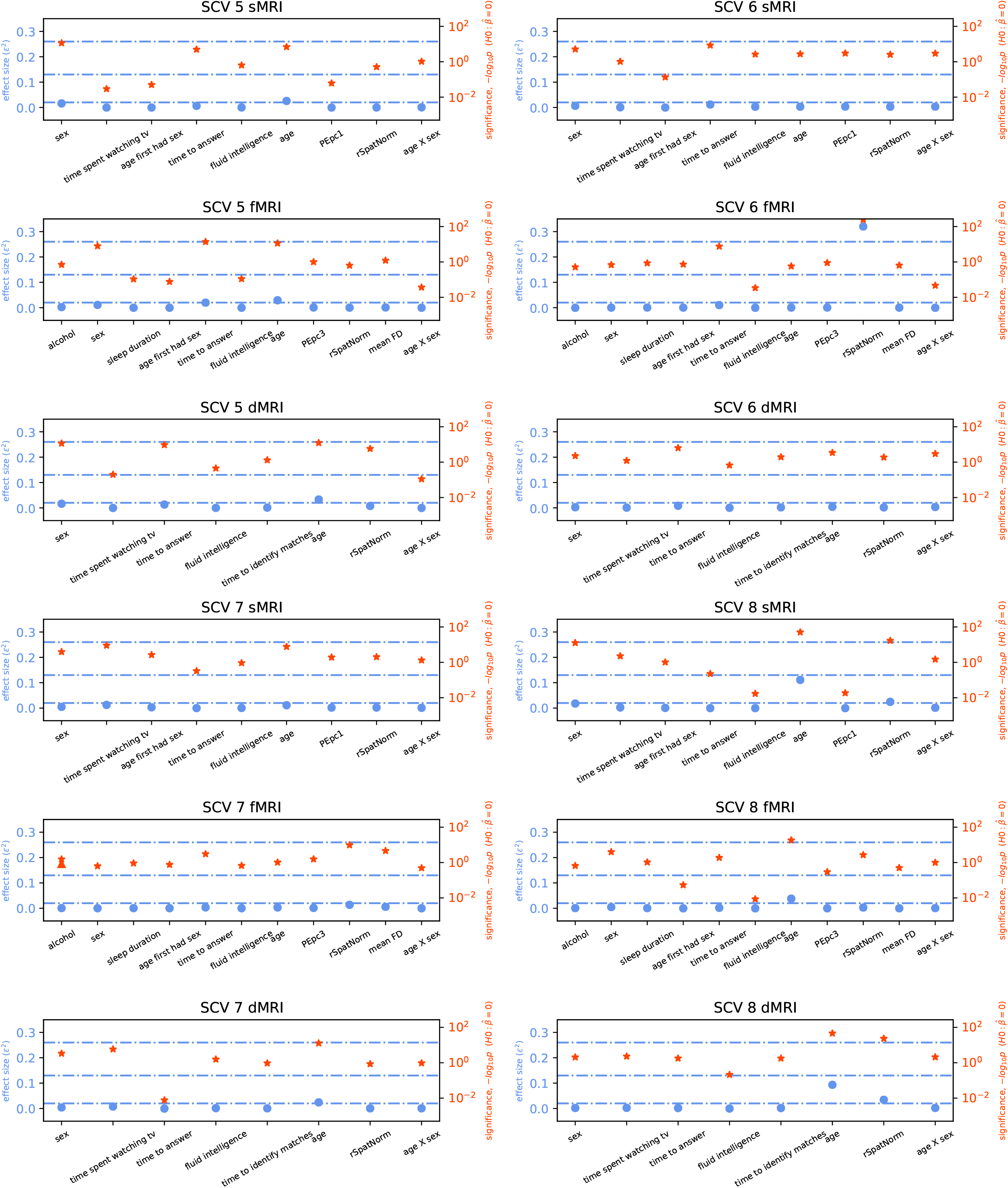

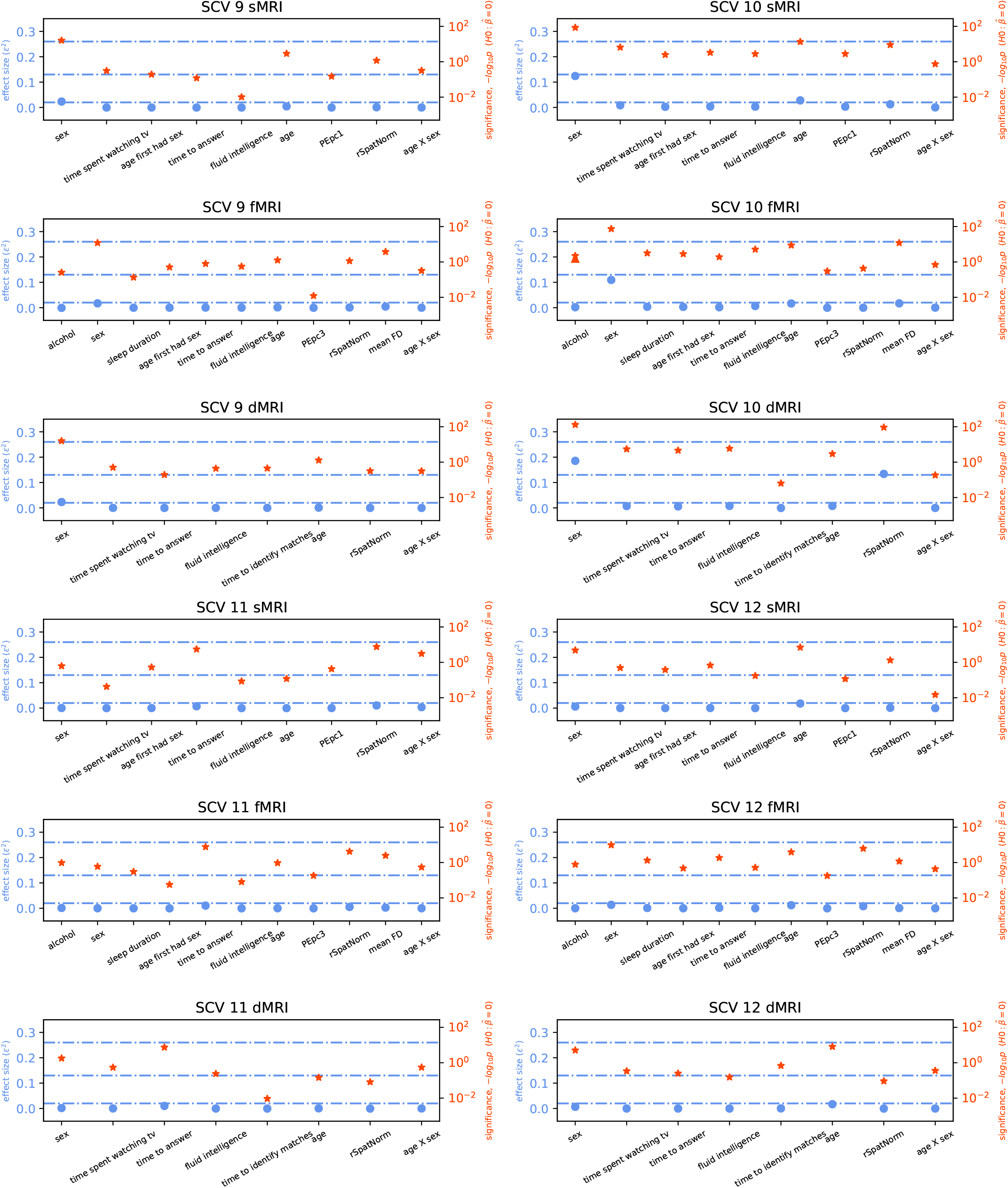

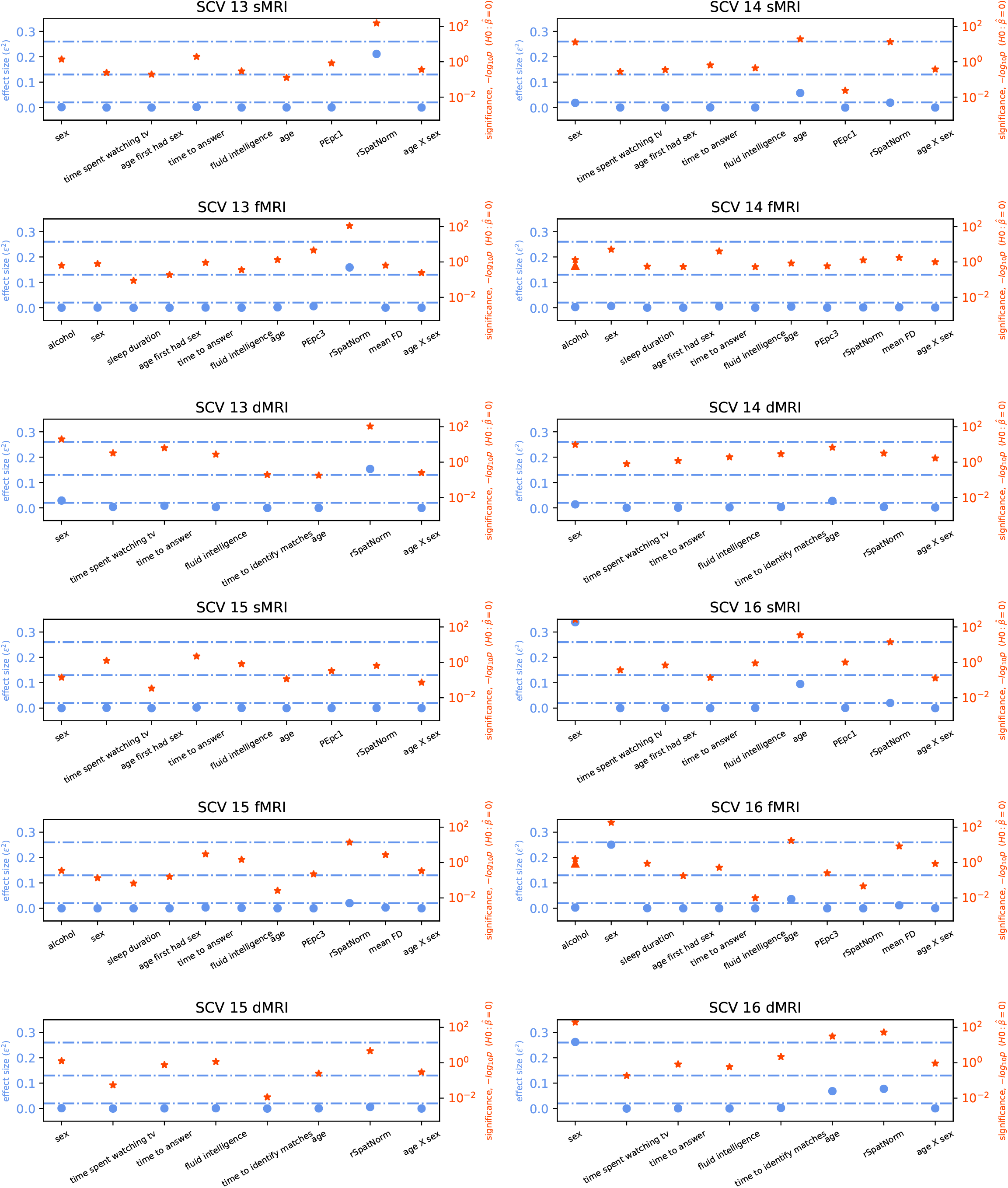

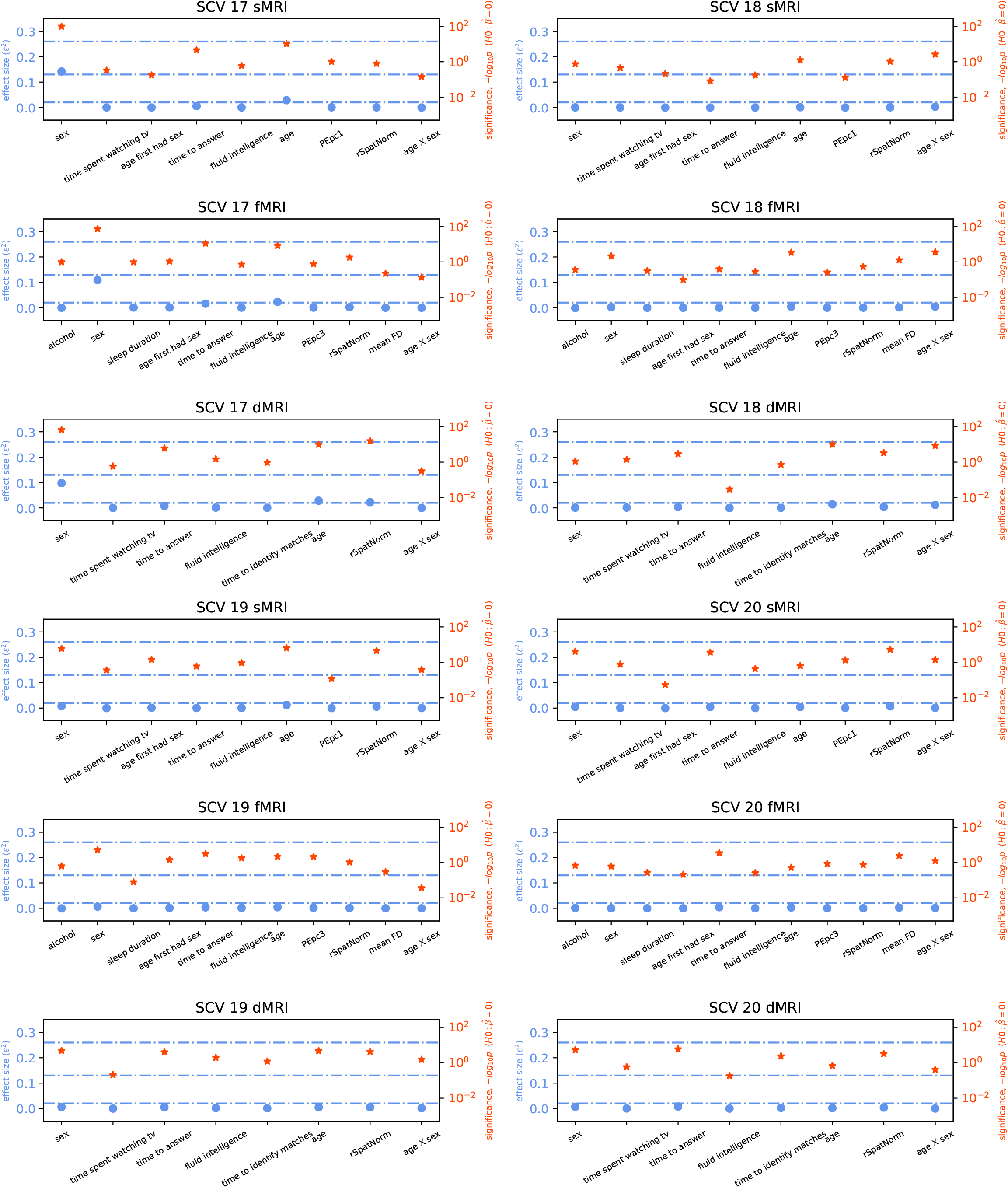

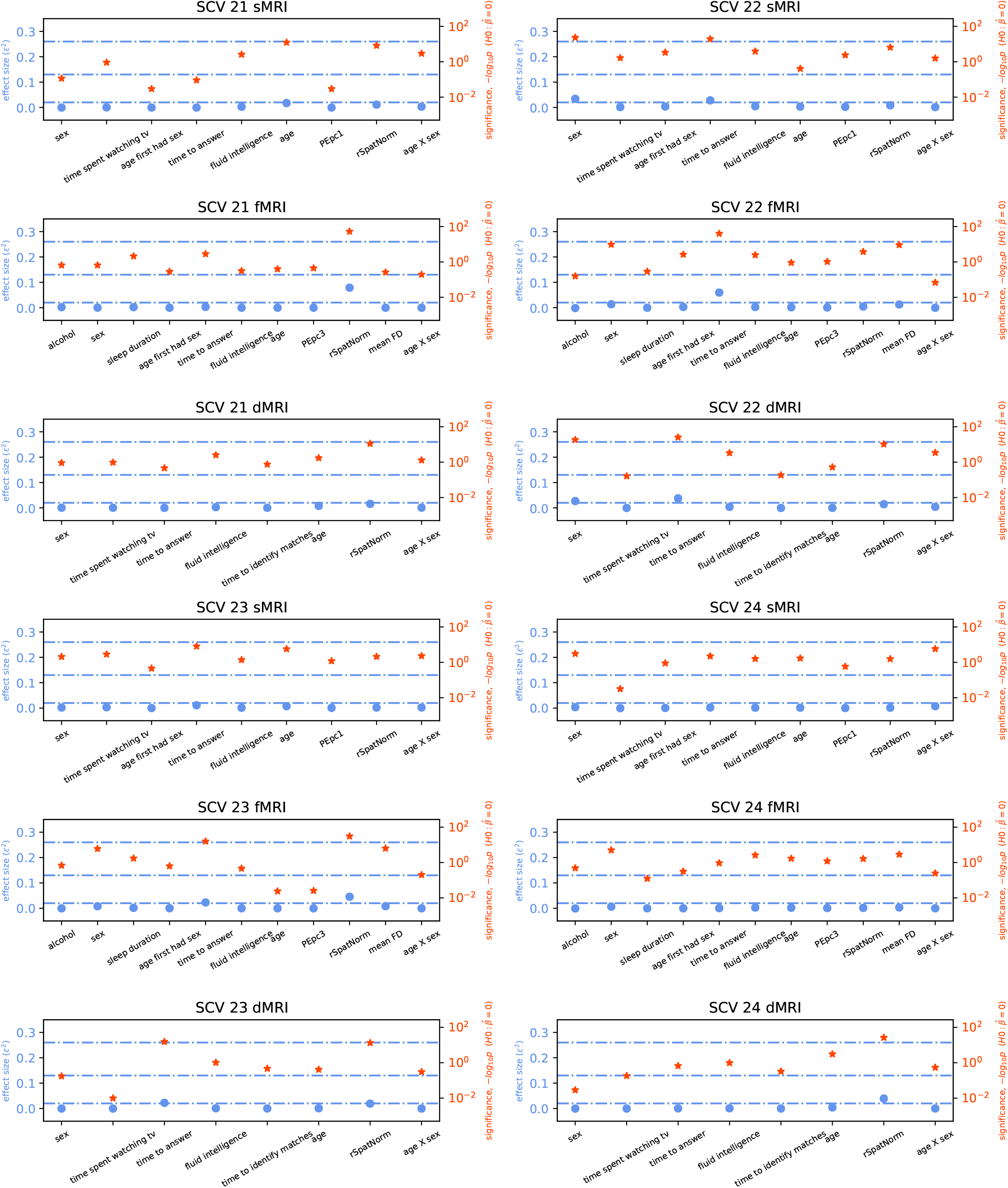

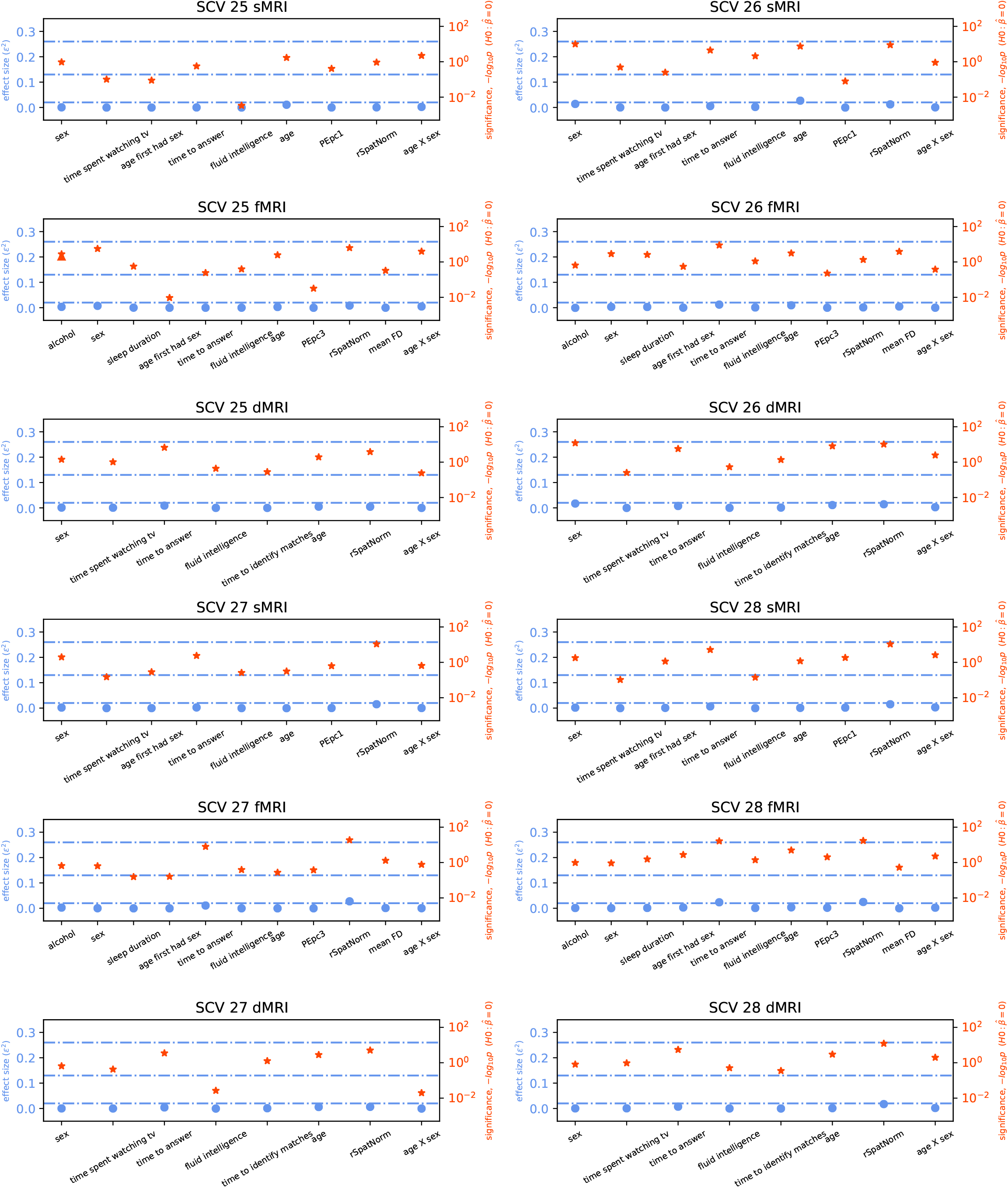

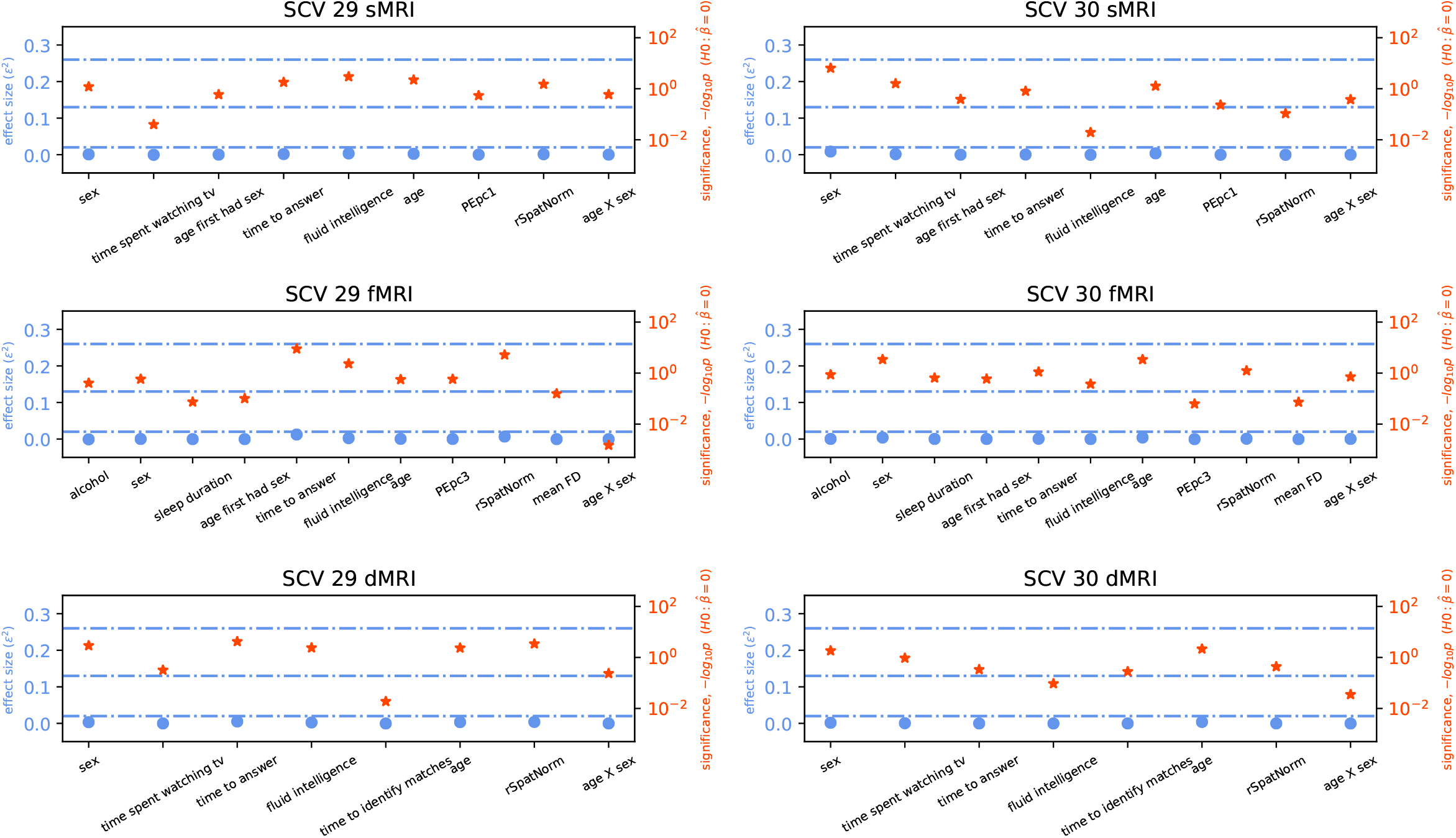

### 6.7. Effect Size Measure (Patient Dataset)

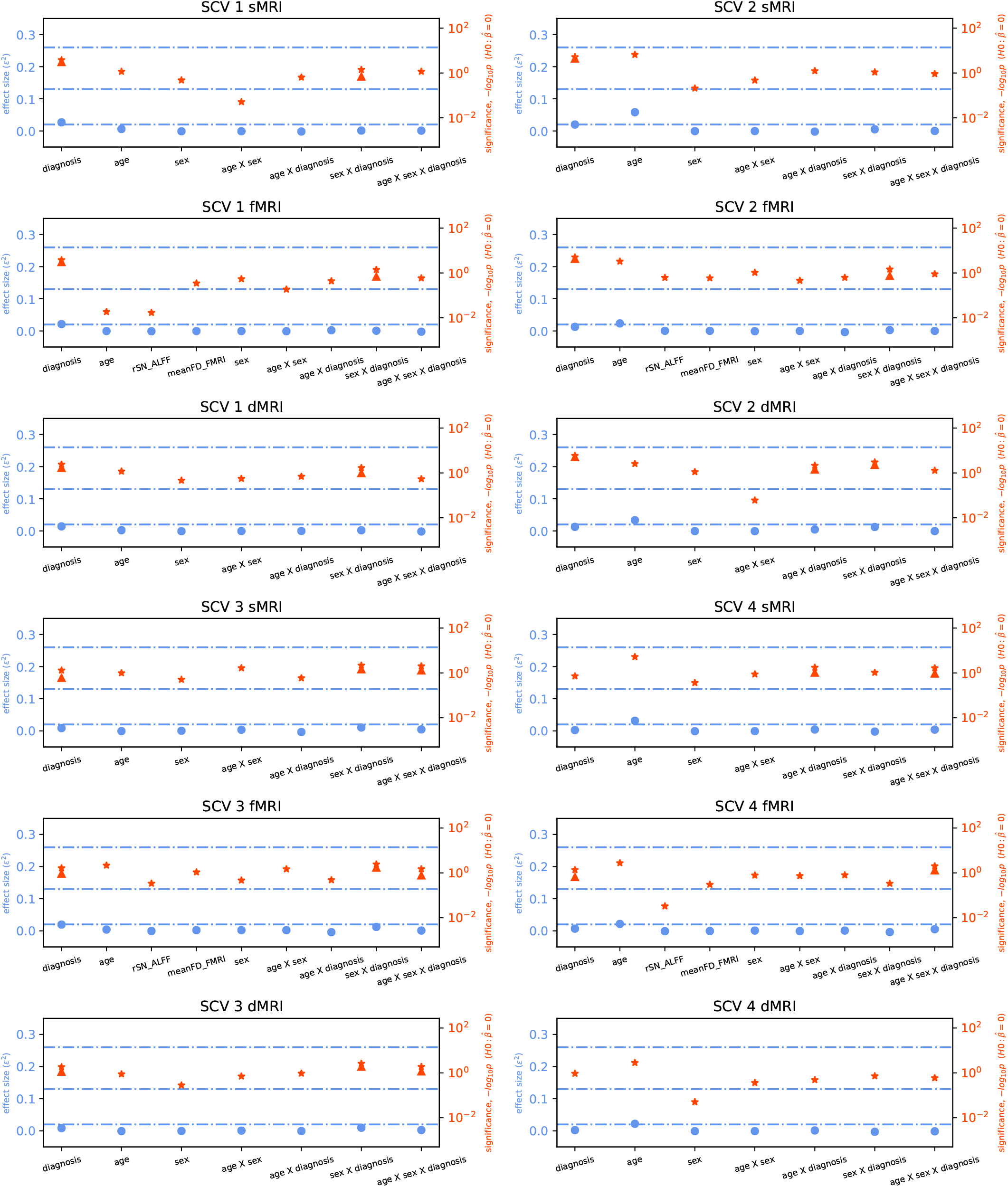

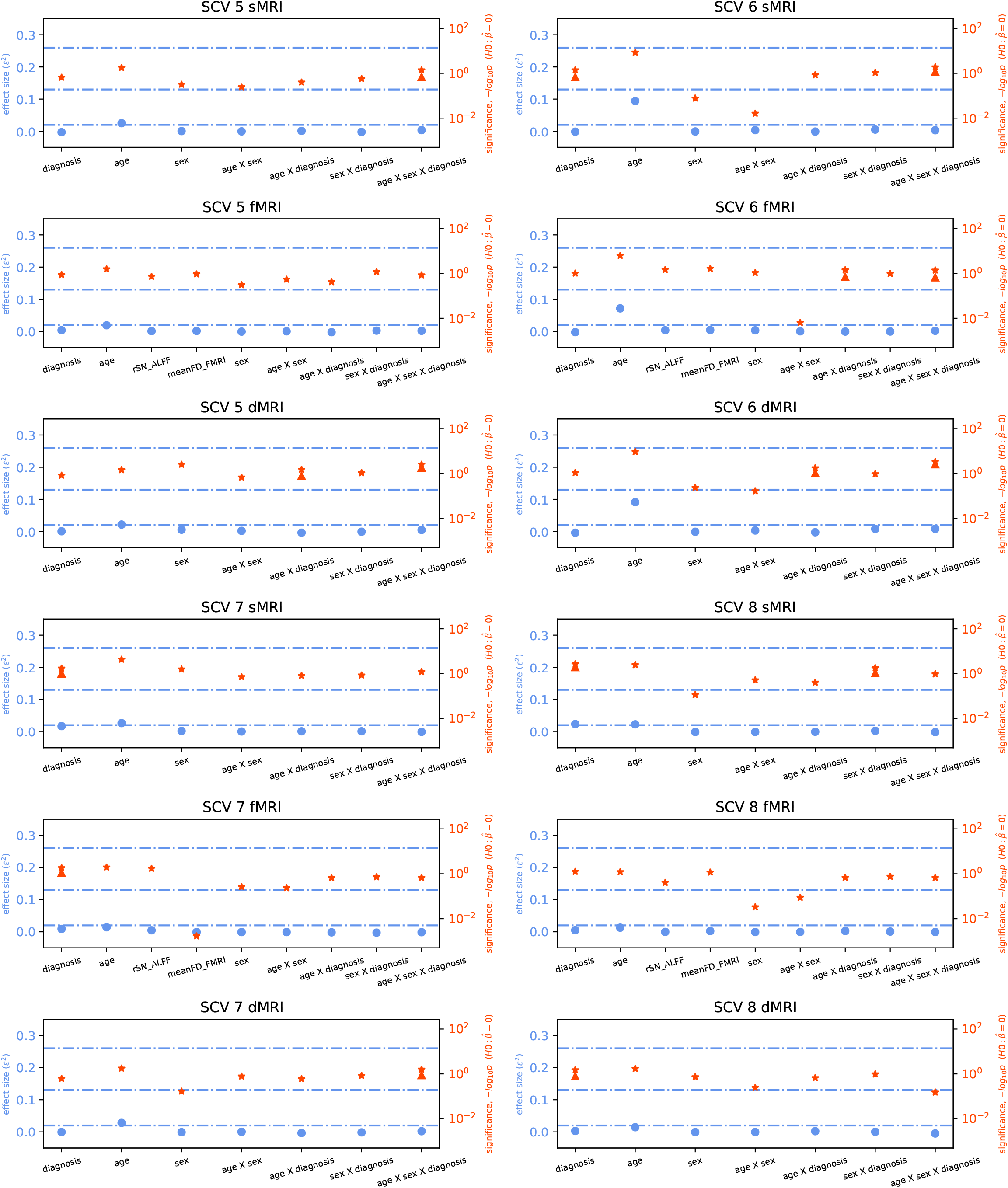

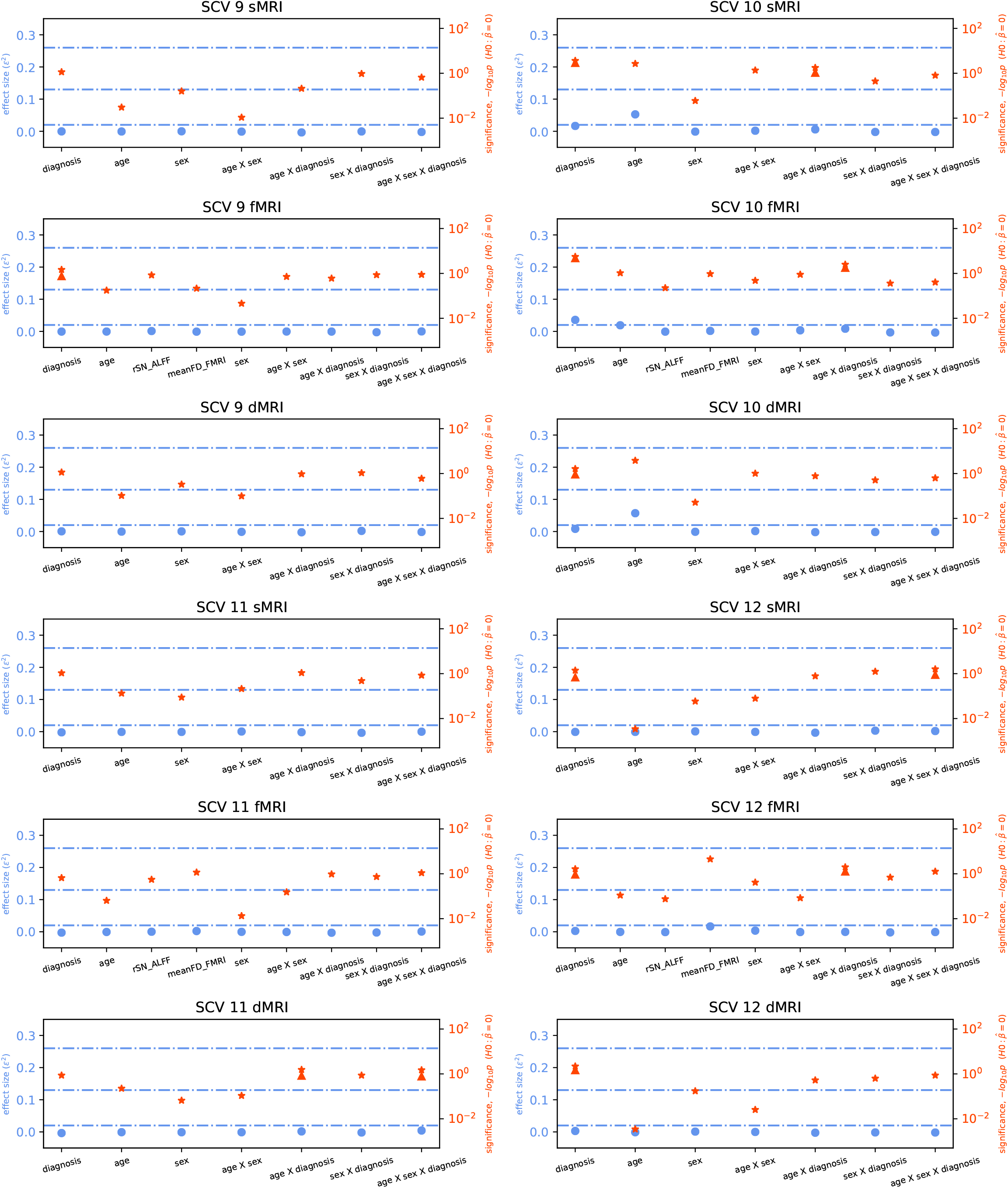

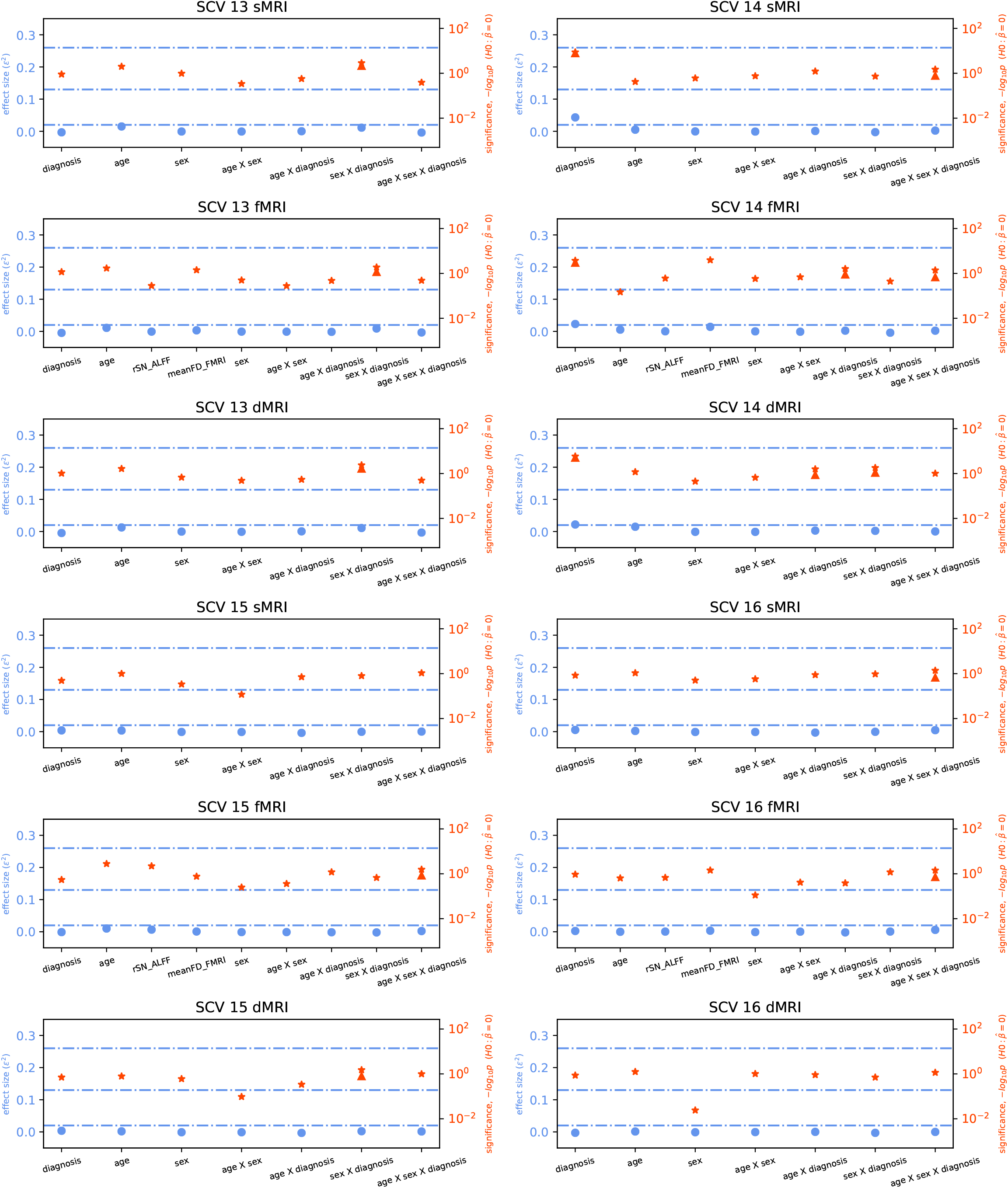

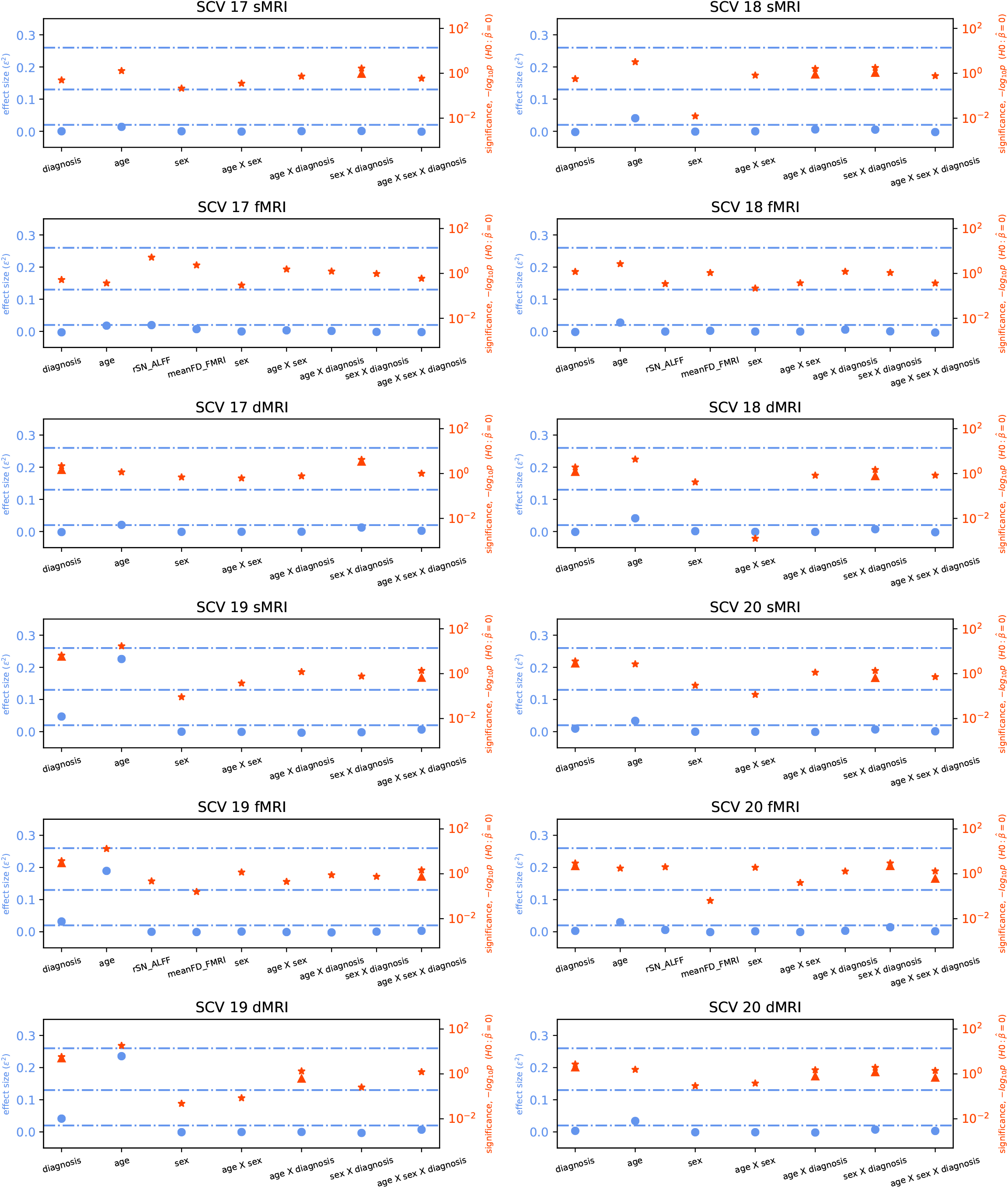

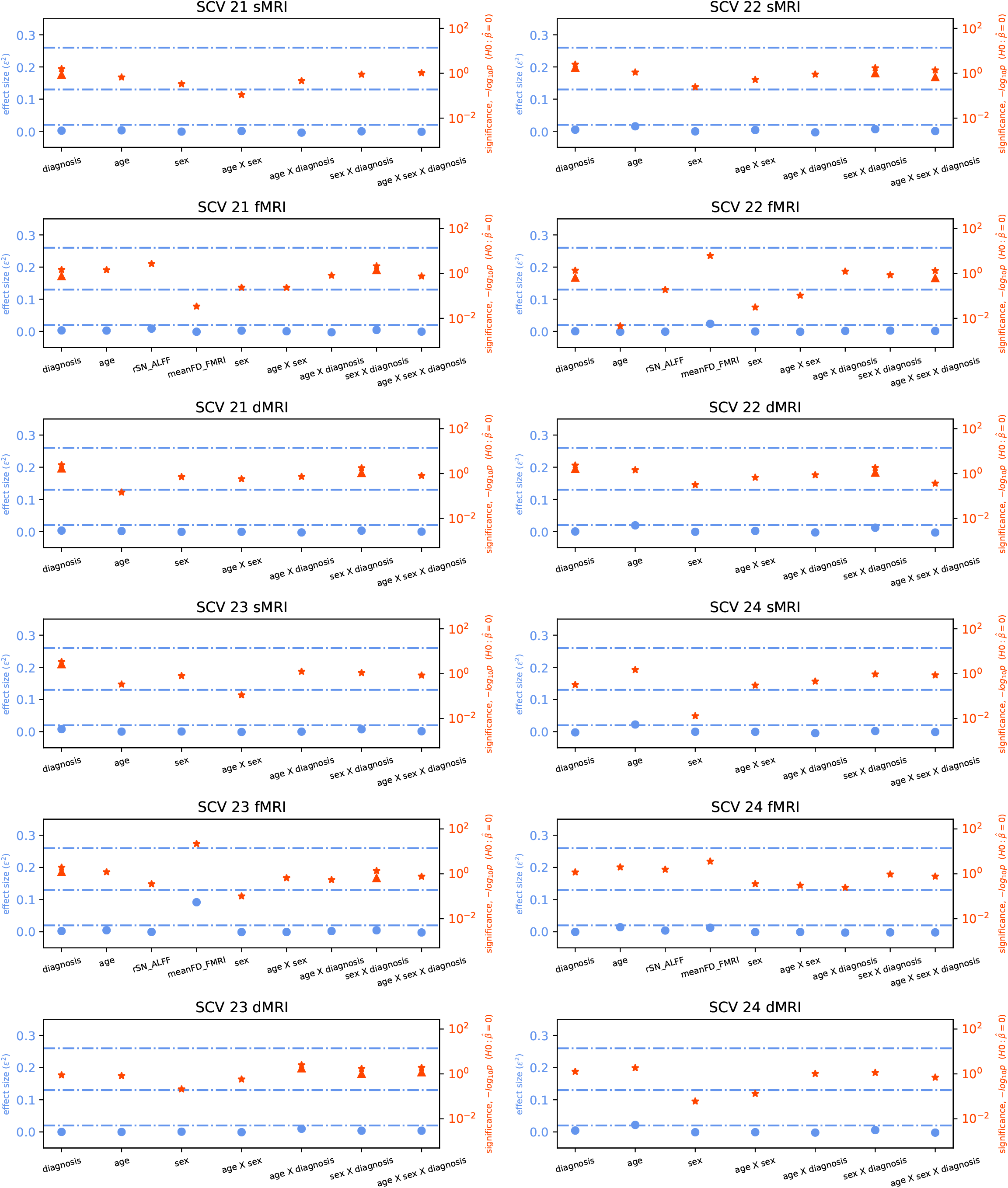

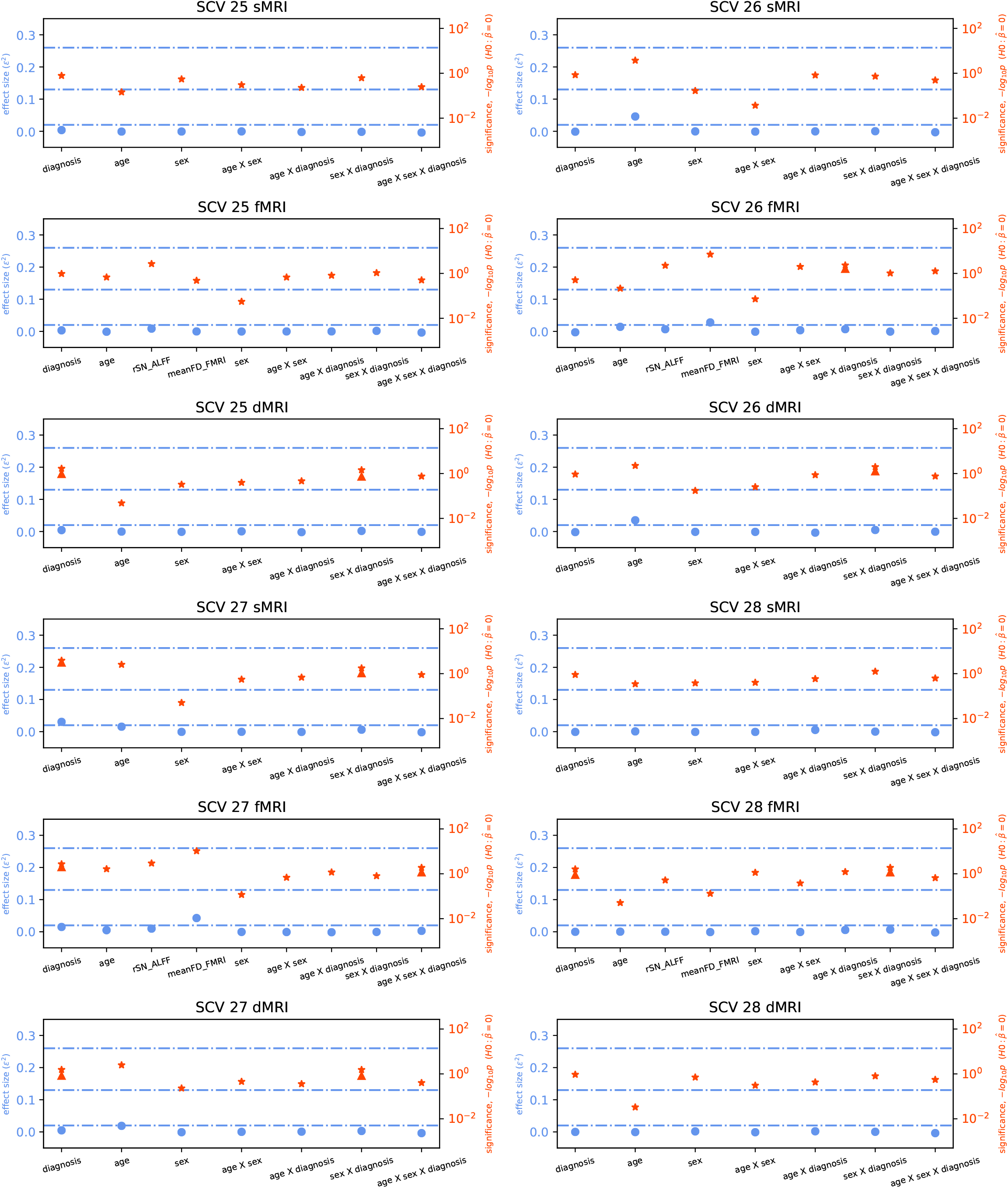

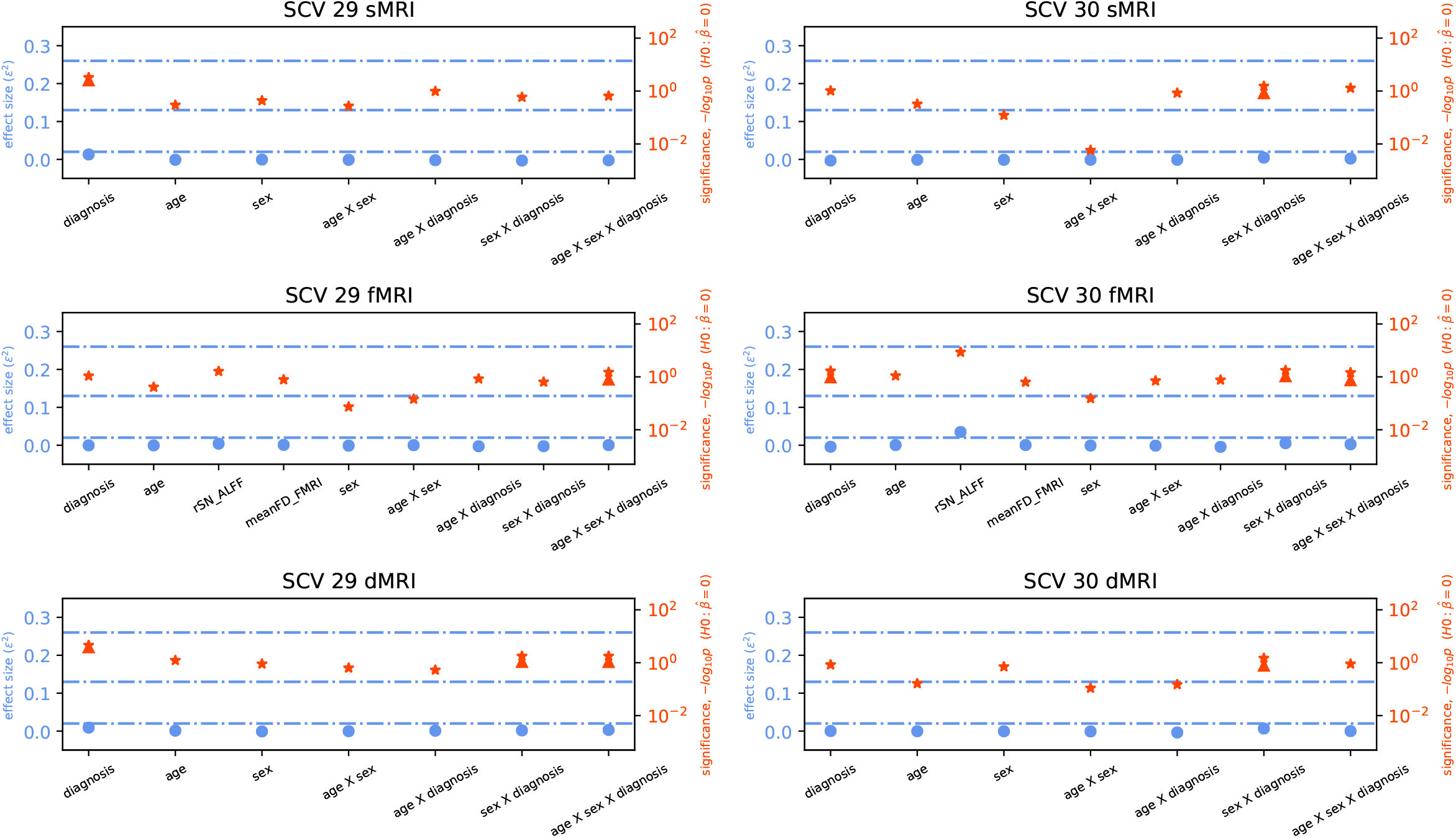

## Footnotes

1 including those due to known and unknown confounds

2 Mean removal is performed before and after partialling. Partialled variances are adjusted to 1.

3 Strictly speaking, we could have regressed out only the sources within the same SCV since all others are independent by the definition of SCVs. Results were nearly identical with such approach.

